# LRH-1 is a novel regulator of neutrophil-driven immune responses within the tumor microenvironment

**DOI:** 10.1101/2025.09.16.676030

**Authors:** Yu Wang, Bryan Duong, Sam Keeley, Natalia Krawczynska, Shruti V. Bendre, Claire P. Schane, Erin Weisser, Nadia Cobo Vuilleumier, Lara Kockaya, Yifan Fei, Anasuya Das Gupta, Hashni Epa Vidana Gamage, Adam T. Nelczyk, Sisi He, Hannah Kim, Jenny Drnevich, Benoit R. Gauthier, Erik R. Nelson

## Abstract

Elevated plasma cholesterol levels have been linked to worse outcomes in breast and ovarian cancer. Prior work including our own has demonstrated that myeloid immune cells are highly responsive to cholesterol fluctuations and to proteins involved in cholesterol regulation. However, the specific roles of Liver Receptor Homolog-1 (LRH-1, or NR5A2), a key transcriptional regulator of cholesterol homeostasis, within myeloid cells remains largely undefined. Interestingly, LRH-1 mRNA levels are reduced in both breast and ovarian tumors compared to normal tissue. Its elevated expression within tumors is associated with increased survival time. These clinical correlations prompted us to explore the role of LRH-1 in myeloid cells particularly in the context of breast and ovarian cancer progression.

Initial analyses confirmed LRH-1 expression in various myeloid cell types, with particularly high levels in neutrophils. We therefore focused on how LRH-1 influences neutrophil behaviors relevant to cancer, including migration, NETosis, phagocytosis, and interactions with T cells. Small molecule ligands for LRH-1 regulated neutrophil migration towards cancer cells. Phagocytosis was also regulated by LRH-1 small molecule ligands, the extent being dependent on type of bait (*e. coli* vs. cancer cells). In T cell co-cultures, neutrophils pretreated with an LRH-1 agonist promoted greater T cell expansion - particularly in CD4⁺ cells - while LRH-1 inhibition suppressed this response. Moreover, LRH-1 activation reduced NETosis, while treatment with an antagonist or inverse agonist enhanced NETosis.

Treatment of mice with an inverse agonist of LRH-1 increased the growth of 4T1 and E0771 mammary tumors. Given prior evidence that neutrophils contribute to the recurrence of dormant lesions, we further examined the role of LRH-1 in this context. Mice harboring dormant D2.0R mammary cancer lesions treated with an LRH-1 inverse agonist exhibited earlier tumor recurrence and enhanced metastatic progression compared to vehicle-treated controls. In contrast, treatment with BL001, a small chemical LRH-1 agonist, delayed recurrence in D2.0R-grafted mice.

Collectively, these findings reveal that LRH-1 is a novel regulator of neutrophil-driven immune responses within the tumor microenvironment. LRH-1 thus remerges as a promising therapeutic target to suppress metastasis or prevent recurrence in breast cancer.

## INTRODUCTION

Despite recent clinical advances, cancers of the breast and ovary continue to have high mortality rates. For breast cancers, associated mortality is due to metastatic recurrence, with approximately 30% of all breast cancer patients experiencing recurrence. For ovarian cancer, the majority of patients are diagnosed at advanced disease typically characterized by peritoneal spread. Unfortunately, immune therapy for breast and ovarian cancers has been disappointing. This is for many reasons, but a major mechanism is thought to be the infiltration of highly immune-suppressive myeloid immune cells (1,2). Thus, new ways to regulate myeloid cells are considered to be the next therapeutic breakthrough in immune based cancer therapies. Interestingly, elevated circulating cholesterol is associated with poor prognosis for both tumor types (3). Similarly, a gain of function germline mutation in PCSK9 drives metastatic progression of breast cancer (4). On the other hand, use of cholesterol lowering drugs (statins) are associated with improved survival (3). Collectively these clinical data suggest that while perhaps not carcinogenic in and of itself, cholesterol influences the progression of breast and ovarian cancers.

There are several mechanisms by which cholesterol influences tumor pathophysiology, recently reviewed in (3). These include the ability to directly regulate downstream signaling molecules such as those in the WNT pathway (5) or anchoring lipid rafts and thus promoting downstream signaling (6–9). Additionally, oxysterol metabolites such as 27-hydroxychoelsterol (27HC) act as ligands for various receptors including the estrogen receptors (ERs) and liver X receptors (LXRs). In ER-positive tumors, 27HC stimulates proliferation of cancer cells (10).

Intriguingly, cholesterol metabolites and regulators of cholesterol homeostasis have been shown to have profound regulatory effects on different immune cell types (11). For example, 27HC works through the LXR to robustly shift myeloid immune cells into an immune suppressive phenotype. After myeloid cells have been exposed to 27HC, they robustly attenuate T cell expansion and activity, thereby promoting breast and ovarian cancer progression (12–15). As cholesterol is catabolized, its bile acid metabolites signal through the farnesoid x receptor (FXR) to upregulate NR0B2; NR0B2 also known as small heterodimer partner (SHP), an atypical nuclear receptor that represses LRH-1 and LXR activity. In myeloid immune cells, NR0B2 suppresses the inflammasome. By doing so, it decreases the ability of these cells to support the expansion of regulatory T cells (T_regs_) – a cell type that is highly immune-suppressive (16,17). In murine models, loss of myeloid cell expression of NR0B2 results in increased tumor growth and metastasis (16). NR0B2 agonists such as DSHN-OMe result in decreased T_regs_ and tumor growth, highlighting NR0B2 and other regulators of cholesterol homeostasis as therapeutic targets (18). Therefore, proteins that are involved in cholesterol homeostasis may represent potential targets to regulate myeloid immune cell function to enhance cancer treatment strategies.

Since 27HC was found to robustly impair myeloid cell mediated anti-cancer immunity through the LXR, this protein presents as a putative therapeutic target. However, the LXR can be selectively modulated, meaning that different ligands have different effects, and those effects are materialized in a context and cell-type specific way (15,19). While the partial agonist 27HC works to shift myeloid cells into a highly immune-suppressive phenotype (14,15) and an inverse agonist stimulates anti-cancer CD8 T cell activity (20), the synthetic LXR agonist, GW3965 reduces myeloid derived suppressor cells (3,21). Although this complexity provides pharmacologic opportunity, more mechanistic insight is needed before we can exploit it as a therapeutic target.

Therefore, we sought to identify alternate targets that might have more near-term impact. Given the clinical associations between cholesterol and cancer, and that we and others have found that myeloid cell functions can be regulated by different aspects of cholesterol homeostasis (3,11), we performed a search for proteins that are (A) associated with survival and (B) expressed in myeloid immune cells. We have previously identified NR0B2 and ABCA1 using a similar approach (16–18,22). This led to the identification of LRH-1 (NR5A2).

Similar to LXR, LRH-1 (NR5A2) is known to upregulate enzymes involved in cholesterol catabolism, such as CYP7A1 and CYP8B1. LRH-1 is a key regulator of cholesterol metabolism but remains poorly understood in the context of immune function and cancer. This is particularly relevant given evidence that LRH-1 exerts context-dependent effects, driving tumor growth in some settings while correlating with improved survival in others. Thus, we performed a series of experiments to elucidate whether LRH-1 influences myeloid cell function, particularly in neutrophils where expression is high, and how this regulation shapes tumor progression. Interestingly, we found that LRH-1 serves to regulate several of their functions, including migration, phagocytosis, support of T cells, and NETosis. The findings reported herein deepen our understanding of LRH-1 in regulating myeloid immune cell function and how it influences cancer progression. Additionally, our findings identify LRH-1 as a potential new therapeutic target to re-shape the tumor-microenvironment, that could potentially reduce the risk of recurrence and enhance the effectiveness of immunotherapy.

## RESULTS

### LRH-1, a known regulator of cholesterol catabolism, is highly expressed in neutrophils – immune cells associated with poor prognosis

Transcriptomic analysis highlighted LRH-1 (NR5A2) as a candidate regulator of myeloid function. In peripheral blood mononuclear cells (PBMC) from healthy individuals, LRH-1 is highly expressed in granulocytes, specifically in basophils and neutrophils (**Fig. 1A**, Monaco dataset, acquired from the Human Protein Atlas (23–25)). Within human breast tumors, single cell RNA sequencing (RNAseq) also indicated increased expression of LRH-1 in myeloid granulocytes particularly neutrophils and mast cells, compared to other immune cells (**Fig. 1B**; GSE114727, accessed by TISCH2 (26,27)). We confirmed that expression was elevated in murine derived neutrophils and the human neutrophil-like HL60 cell line compared to other myeloid cell types or mammary cancer cells (**Supplementary Fig. 1A-B**). Using the GEPIA2 platform (28), we found that LRH-1 was significantly decreased in tumor tissue compared to normal tissue in several cancer types, especially for breast and gynecologic tumors (ovarian, cervical squamous cell carcinoma, uterine corpus endometrial carcinoma and uterine carcinosarcoma) (**Fig. 1C**).

**Figure 1.**
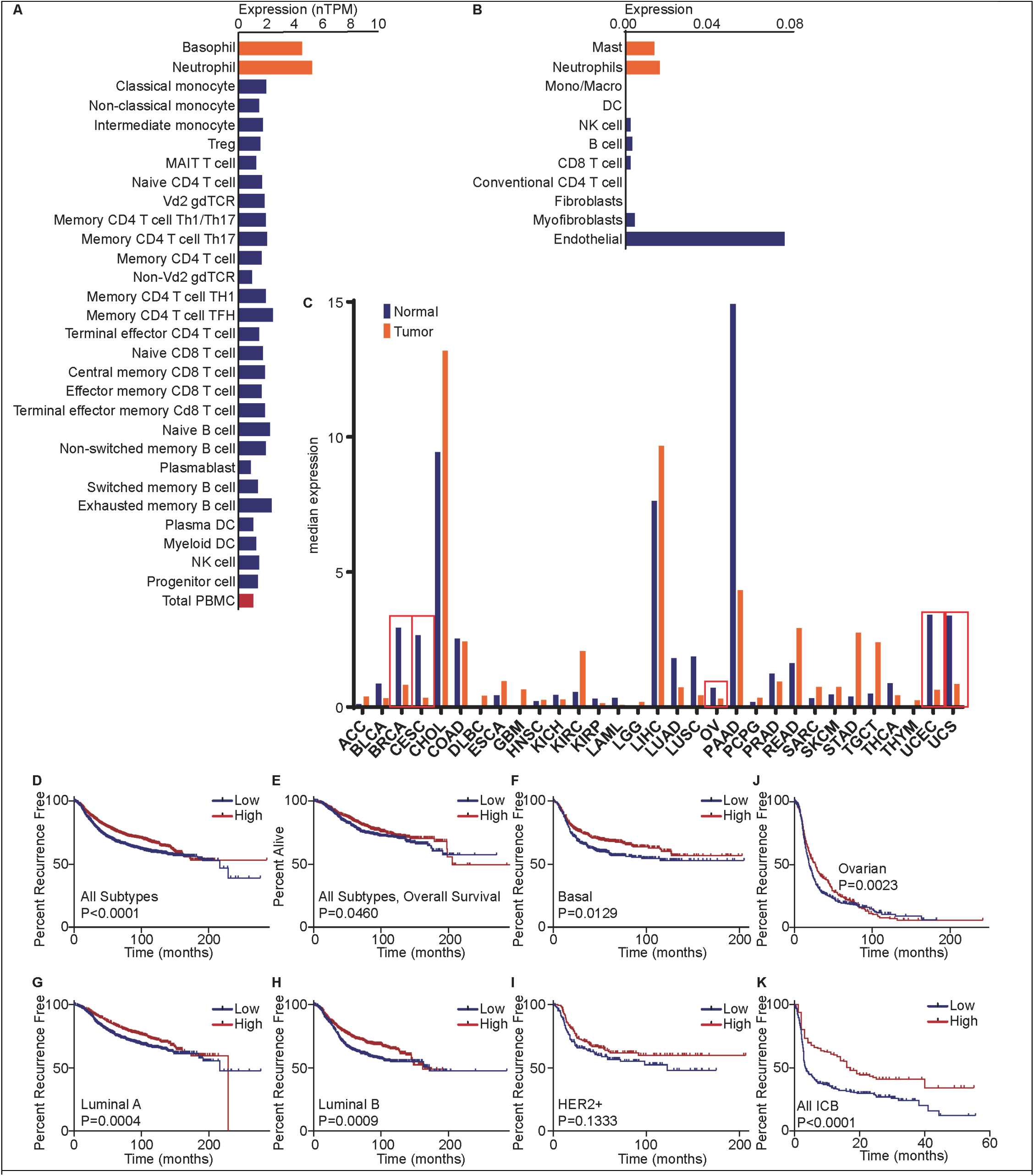
LRH-1 is expressed in granulocytes within mammary tissue and tumors, is decreased in mammary tumors compared to normal, and its expression in breast tumors is associated with increased survival. (**A**) Expression of *LRH-1* in different peripheral blood mononuclear cells (PBMCs) isolated from humans (dataset from (25), accessed through the Human Protein Atlas (23,24)). (**B**) RNA expression of *LRH1* in annotated cell types through single cell RNAseq. Data from GSE114727, extracted from TISCH2 (26,27)). (**C**) RNA expression of *LRH-1* in healthy versus tumor tissue. Data extracted from GEPIA2 (28). (**D**) Elevated *LRH-1* expression in breast tumors is associated with increased recurrence free survival. All breast cancer subtypes are considered here. (**E**) Elevated *LRH-1* expression in breast tumors is associated with increased overall survival. All breast cancer subtypes are considered here. (**F**) Elevated *LRH-1* expression in Basal type breast tumors is associated with increased recurrence free survival. (**G**) Elevated *LRH-1* expression in Luminal A type breast tumors is associated with increased recurrence free survival. (**H**) Elevated *LRH-1* expression in Luminal B type breast tumors is associated with increased recurrence free survival. (**I**) *LRH-1* expression in HER2+ type breast tumors is not significantly associated with recurrence free survival (P=0.1333). (**J**) Elevated *LRH-1* expression in ovarian tumors is associated with increased progression free survival. (**K**) Elevated *LRH-1* expression in tumors is associated with response to immune checkpoint blockade (ICB). All tumor types included in the database were used in this analysis. (**D-K**) Data was obtained from the Kaplan-Meier Plotter webtool, which pools data from GEO, EGA, and TCGA (36). *LRH-1* expression in tumors was parsed by the median except for (J). The P value reported is from testing by Log-rank (Mantel-Cox method), except for (J) where the Gehan-Breslow-Wilcoxon test was used.

When assessing breast tumors, we found that elevated LRH-1 expression was associated with increased recurrence free and overall survival (**Fig. 1D-E**, data from the Kaplan-Meier Plotter (29), which populates data from GEO, EGA, and TCGA). Stratification by subtype, elevated LRH-1 expression was associated with improved recurrence free survival for the basal, luminal A and luminal B tumors (**Fig. 1F-I**). Similarly, increased LRH-1 expression in tumors was associated with improved progression free survival in ovarian cancer (**Fig. 1J**).

Interestingly, LRH-1 expression is associated with a robust improvement in response to immune checkpoint blockade (ICB) (**Fig. 1K**). For this analysis all tumor types were considered, since there is very little transcriptomic data publicly available for ICB-treated breast cancer tumors. These clinical associations with survival across different breast cancer subtypes, ovarian cancer and those undergoing ICB therapy suggest that LRH-1 expression in a cell type such as neutrophils common to all tumors, is mediating its beneficial effects.

### Small molecule LRH-1 modulators differentially affect proliferation of mammary and ovarian cancer cell lines, in vitro

In order to assess potential cancer cell-intrinsic effects of LRH-1, we evaluated the effects of small molecule modulators on cancer cell proliferation. Since expression of LRH-1 was high in neutrophils, we focused on murine syngeneic models in anticipation of evaluating *in vivo* tumor growth in immune-competent models. Treatment with the agonist, RR-RJW100 had no significant effect on proliferation in either ovarian ID8 cancer cells or mammary cancer D2.0R cells, but modestly reduced the proliferation of mammary cancer 4T1 and E0771 cells (**Fig. 2**). An LRH-1 antagonist (CAS #1185410-60-9 named “LRH-1 antagonist”) had no effect on ID8 cells, but decreased proliferation of 4T1, D2.0R and E0771 cells. On the other hand, SR-1848 which is described as an LRH-1 inverse agonist, significantly attenuated proliferation across all four cell lines tested (**Fig. 2**). The effects of SR1848 were similar to several other reports indicating that LRH-1 activity drives proliferation (3). However, given the clinical data that tumoral LRH-1 expression is associated with increased survival, these findings were counter-intuitive. Thus, we reasoned that immune cell-mediated effects, dominate over tumor intrinsic response *in vivo*.

**Figure 2.**
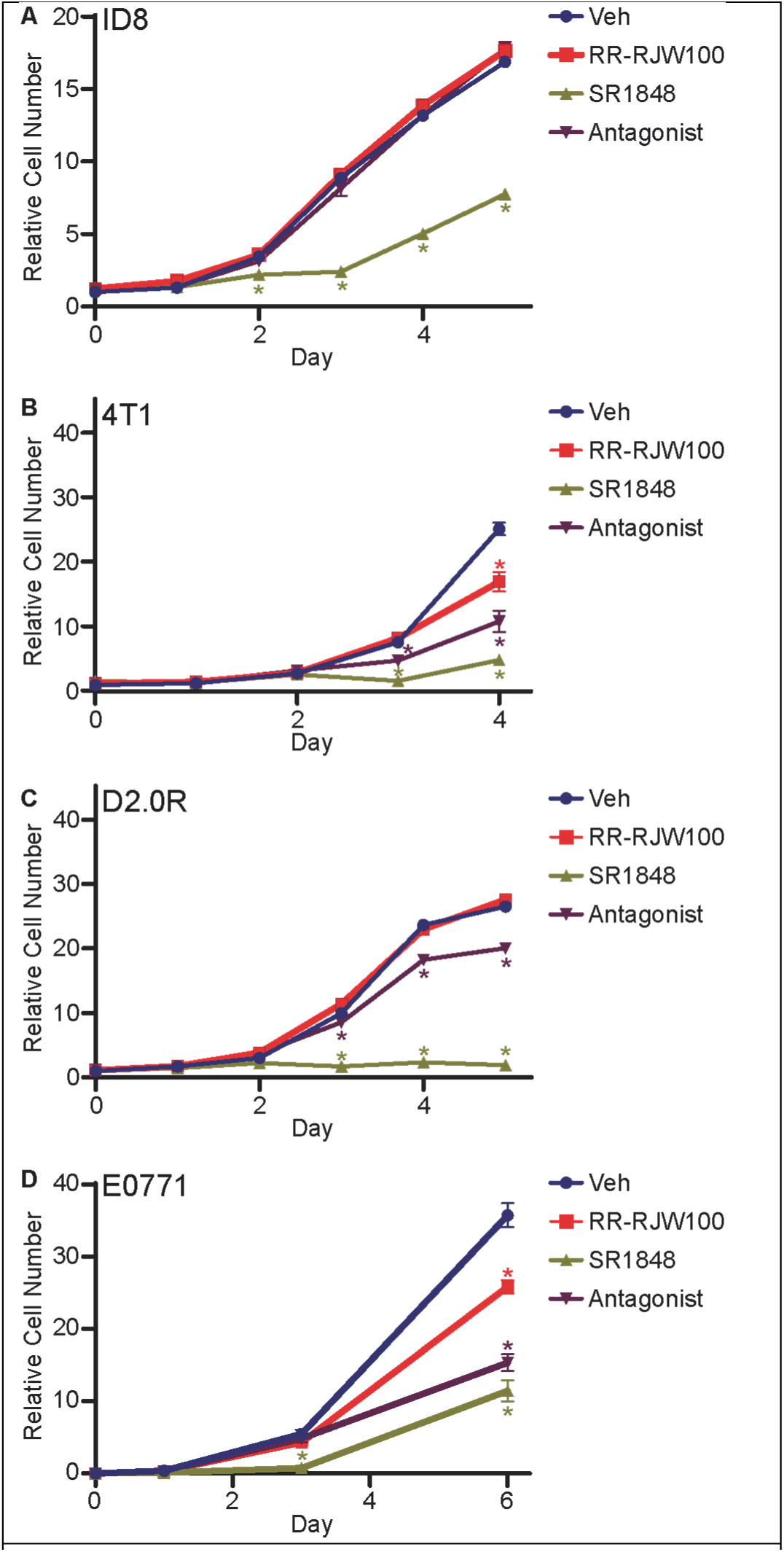
LRH-1 inverse agonist robustly decreases proliferation of ID8 ovarian cancer cells, and 4T1, D2.0R and E0771 mammary cancer cells. Cells were plated and treated with indicated ligands every other day. At indicated time-points, plates were frozen and total DNA quantified as a proxy for cell number, using Hoescht staining (10). Different colored asterisks indicate statistically significant differences between that treatment group and vehicle (P<0.05, two-way repeated measures ANOVA followed by multiple comparison test with Šidáks correction). (**A**) ID8 ovarian cancer cells. (**B**) 4T1 mammary cancer cells. (**C**) D2.0R mammary cancer cells. (**D**) E0771 mammary cancer cells.

### Small molecule modulators of LRH-1 alter expression of genes implicated in neutrophil function

Given that neutrophils had high expression of LRH-1 and that neutrophils (or granulocytic myeloid derived suppressor cells; GMDSCs) are abundant in breast tumors and their metastatic lesions (30), we evaluated how LRH-1 might regulate functions known to influence tumor pathophysiology.

Neutrophils are often short-lived, first responders. They are generalized phagocytes that secrete hydrogen peroxide, hypohalous acids, nitric oxide, cytokines etc. (31). They can also undergo a process termed NETosis, where they extrude their DNA and associated proteins to form NETs that can encapsulate pathogens or foreign particles (32). Due to the volatile nature of neutrophils, we have relied on several known small molecule modulators of LRH-1 function (**Supplementary Fig. 2A**). To determine the optimal dose to use in neutrophils, we considered their EC/IC50s and performed a dose response to assess viability (**Supplementary Fig. 2B**). We then used these small molecule modulators to probe how LRH-1 activity influences different neutrophil functions important for tumor biology.

In order to elucidate the overall effects of LRH-1 activity in neutrophils, bulk RNAseq was performed on murine bone marrow derived neutrophils treated with vehicle, RR-RJW100, LRH-1 antagonist or SR1848. Each replicate was from neutrophils isolated from a different mouse. Treatment with SR1848 resulted in extreme changes in gene expression as well as an apparent batch effect. The changes were so extensive that the overall total RNA pool was substantially shifted, causing nearly all genes to appear “differentially expressed” when applying conventional thresholds. Many of these apparent changes are unlikely to be genuine, but instead reflect this global shift in the RNA pool. To address this, we applied alternative statistical approaches aimed at detecting only the most robust and consistent gene-level changes. Including SR1848 in the same statistical framework as RR-RJW100 and the antagonist would have obscured the comparatively subtle effects of the latter two treatments. Therefore, downstream informatics were performed comparing vehicle to RR-RJW100 or LRH-1 antagonist, and separately for vehicle to SR1848. Heatmaps indicating differentially expressed genes (DEGs) compared to vehicle are shown in **Fig. 3A-B** (differential expression was assessed using empirical Bayes moderated statistics with FDR<0.1 for comparisons between vehicle and RR-RJW100 or LRH-1 antagonist, and the *treat* function with FDR<0.005 for comparisons between vehicle and SR1848). Multidimensional scaling (MDS) plots showed separation of DEGs from vehicle, with dimension 2 revealing differences between Veh and either RR-RJW100 or LRH-1 antagonist (**Fig. 3C**). Interestingly, RR-RJW100 and LRH-1 antagonist separated DEGs to the same direction in dimension 2, although LRH-1 antagonist generally had a larger change. MDS analysis also showed changes between Veh and SR1848, with one batch showing separation in dimensions 1 and 2, and the other showing most separation in dimension 1 (**Fig. 3D**). Because we were interested in identifying pathways potentially regulated by these ligands, we used batch 2 for subsequent analyses. DEGs between the different treatment groups showed a mix of uniqueness as well as overlapping with other ligand treatments (**Fig. 3E**). Compared to vehicle, treatment with the agonist RR-RJW100 had the fewest DEGs compared to the other ligands. Interestingly, RR-RJW100 shared several DEGs with the LRH-1 antagonist and inverse agonist (SR1848) (**Fig. 3E**).

**Figure 3.**
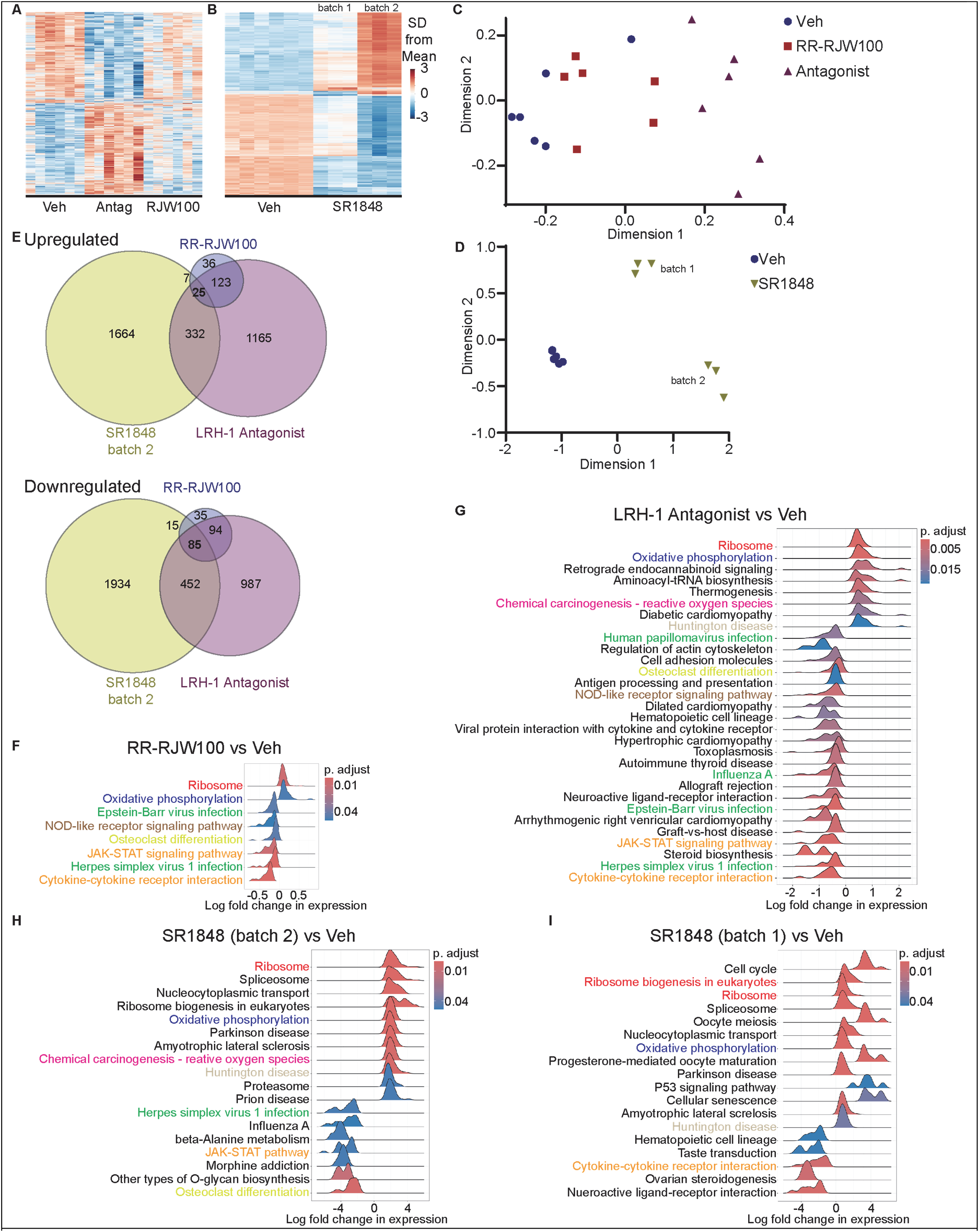
Small molecule modulators of LRH-1 result in unique transcriptional responses in neutrophils. Murine bone marrow derived neutrophils were treated with vehicle, RR-RJW100, LRH-1 antagonist or SR1848. Each replicate is derived from neutrophils from an individual mouse. After 24h of exposure to the indicated ligand, RNA was isolated and subjected to RNA-seq. SR1848 resulted in robust changes in gene expression, and thus was analyzed seperately. (**A-B**) Heatmaps indicating differentially expressed genes (DEGs; for Veh vs. RR-RJW100 or vs. LRH-1 antagonist: FDR<0.1; for Veh vs. SR1848, FDR<0.005. SR1848 treatment showed a potential batch effect; the two batches denoted here). (**C-D**) Multidimensional scaling (MDS) plots. (**E**) Venn diagrams indicating DEGs compared to vehicle for each treatment (upregulated genes are depicted above downregulated genes) (Venn diagrams are adapted from https://bioinforx.com/apps/venn.php). (**F-I**) Kyoto Encyclopedia of Genes and Genomes (KEGG) analysis comparing vehicle to each treatment. Analysis for SR848 was done for each batch, with focus being placed on batch 2. Pathways similar to the different groups are highlighted in colored text. Gene ontology and gene set enrichment analyses are in **Supplementary** Fig. 3.

Pathway analysis confirmed this pattern. Kyoto Encyclopedia of Genes and Genomes (KEGG) analysis comparing vehicle to each treatment revealed several common pathways, including ribosome, oxidative phosphorylation, cytokine / cytokine receptor signaling, viral infection and osteoclast differentiation (**Fig. 3F-I**). Interestingly, most of the enriched pathways were regulated in the same direction (up or downregulation) regardless of ligand. Gene ontology (GO) analysis also revealed several pathways commonly regulated by LRH-1 ligands (**Supplementary Fig. 3A**). When DEGs specific to RR-RJW100 were probed, GO terms associated with cholesterol, lipid and phospholipid transport were enriched (**Supplementary Fig. 3B**). When only those DEGs in common between SR1848 and the LRH-1 antagonist were probed several functional pathways had decreased enrichment, including response to virus, phagocytic vesicle and MHC class 1 receptor activity (**Supplementary Fig. 3C**).

Gene set enrichment analysis (GSEA) further emphasized functional differences, and was performed focusing on neutrophil-specific functions including the complement system, oxidative phosphorylation and apoptosis (**Supplementary Fig. 3D**). The interplay between complement signaling and apoptotic regulation shapes neutrophil functional outcomes in inflammatory environments (33). In our dataset, neutrophils treated with RR-RJW100 showed enrichment of the HALLMARK_COMPLEMENT pathway (NES=1.49, padj=0.022), while also significantly downregulating HALLMARK_APOPTOSIS (NES=-1.61, padj=0.015). In contrast, both LRH-1 antagonist and SR1848 showed no significant enrichment in complement activity and only mild, non-significant suppression of apoptosis. These findings suggest that RR-RJW100 treatment promotes a functional neutrophil phenotype characterized by enhanced survival and immune engagement. Complement activation can augment neutrophil phagocytosis, degranulation, and cytokine release, while reduced apoptotic signaling may prolong neutrophil lifespan, allowing for extended effector activity (34). This phenotype likely supports immune surveillance, clearance of pathogens or debris, and potentially enhanced antigen presentation or T cell priming. In contrast, the absence of complement activation and only partial apoptosis suppression in the LRH-1 antagonist or SR1848 treated neutrophils may shift their cell fate towards alternative functions such as NETosis, rather than sustained immune interactions.

NETosis, particularly mitochondrial NETosis, is tightly linked to increased oxidative phosphorylation (OXPHOS) (35,36). We found that OXPHOS gene sets were significantly enriched in neutrophils treated with either LRH-1 antagonist (NES=1.91, padj=4.2e–6) or SR1848 (NES=1.96, padj=1.2e–6). These results suggest that antagonism or inverse agonism of LRH-1 reprogram neutrophil metabolism toward mitochondrial respiration, a prerequisite for NETosis. The upregulation of OXPHOS is needed to support chromatin de-condensation and NET release, especially under inflammatory conditions (36,37).

### LRH-1 modestly reduces neutrophil migration

Neutrophil infiltration of breast or ovarian tumors is associated with risk of onset (38,39), higher grade (40) and poor prognosis (41–54), especially when considering the neutrophil to lymphocyte ratio. Thus, we next explored whether LRH-1 signaling influenced neutrophil migration towards media that had been conditioned by 4T1 mammary cancer cells (experimental setup depicted in **Fig. 4A**). The LRH-1 agonist RR-RJW100 modestly reduced migration in a dose dependent manner (**Fig. 4B**). The LRH-1 antagonist increased migration (**Fig. 4B**). Interestingly though, the inverse agonist, SR1848, inhibited migration, similar to RR-RJW100 (**Fig. 4B**). These results indicate that LRH-1 signaling regulates migration in a ligand-dependent manner; RR-RJW100 and SR1848 reduce chemotaxis, whereas the antagonist enhances it.

**Figure 4.**
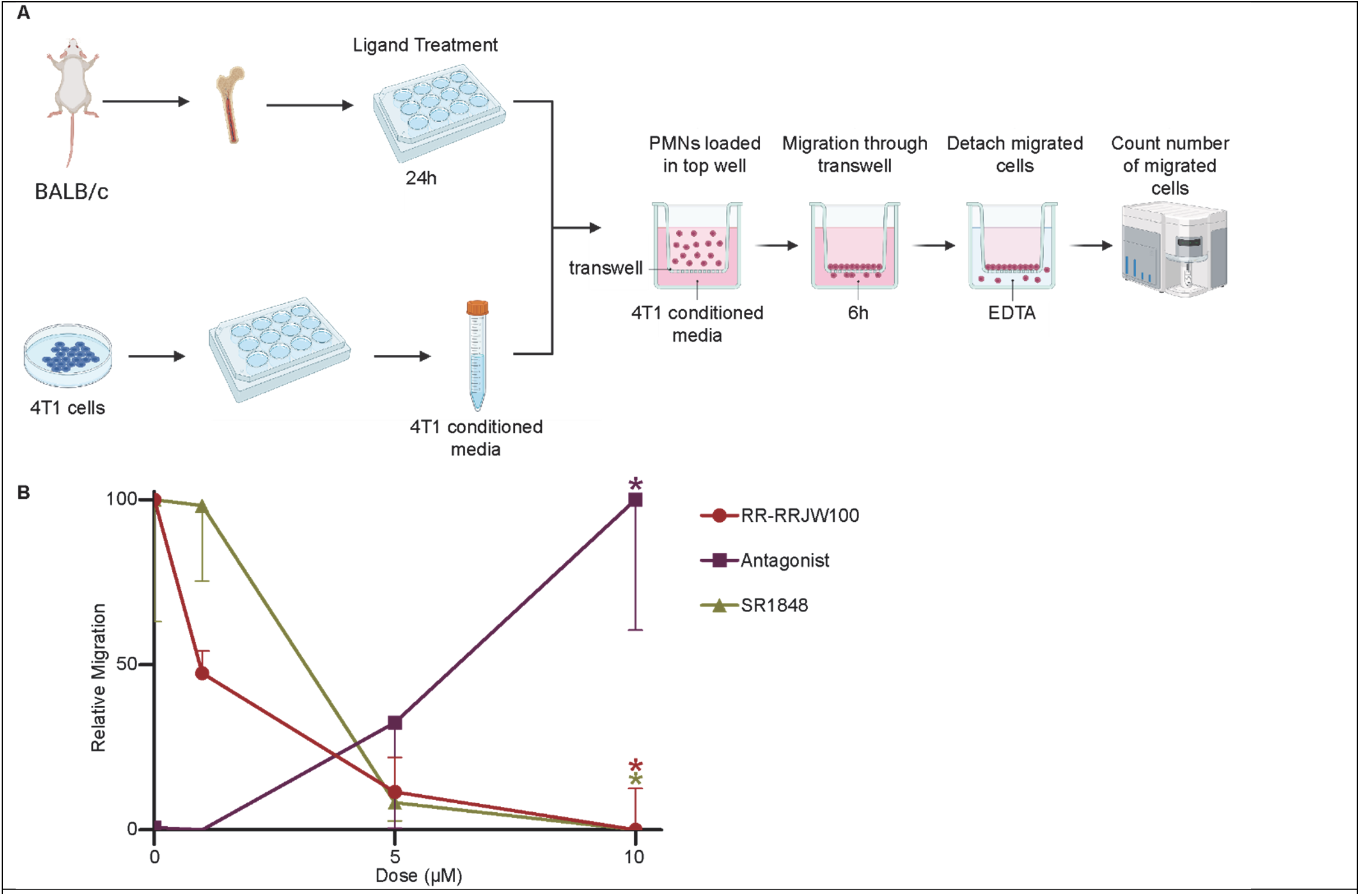
LRH-1 modulators result in altered neutrophil migration towards cancer cell conditioned media. (**A**) Illustration of experimental setup. Neutrophils were isolated from the bone marrow of female BALB/c mice and treated with indicated ligands for 24h. At this point, they were seeded on the top of a Boydon transwell chamber and allowed to migrate towards media that had been conditioned with 4T1 mammary cancer cells. Neutrophils that had migrated through the transwell membrane were quantified by flow cytometry. Adapted from BioRender. (**B**) Quantified data. Data for each treatment were normalized prior to plotting. One-way ANOVA was performed for each treatment with respectively colored asterisks denoting significant differences to vehicle treated (P<0.05, multiple comparison test with Šidáks correction; error bars are shown in one direction for visual clarity).

### Basal LRH-1 activity maintains phagocytosis in neutrophils

A primary task of neutrophils is to perform generalized phagocytosis. To evaluate whether LRH-1 was involved in this process, neutrophils were treated with different LRH-1 ligands prior to assessment of their capacity to phagocytose pHrodo *E. coli* conjugated bioparticles (experimental method outlined in **Fig. 5A**). Treatment with the LRH-1 endogenous agonist, DLPC, significantly impaired phagocytosis, as did the synthetic agonist RR-RJW100 (**Fig. 5B**). The LRH-1 antagonist had no significant effect on phagocytosis. However, the inverse agonist, SR1848 also reduced phagocytosis. Thus, basal LRH-1 activity also maintains phagocytosis, with either loss or gain of LRH-1 activity decreasing it.

**Figure 5.**
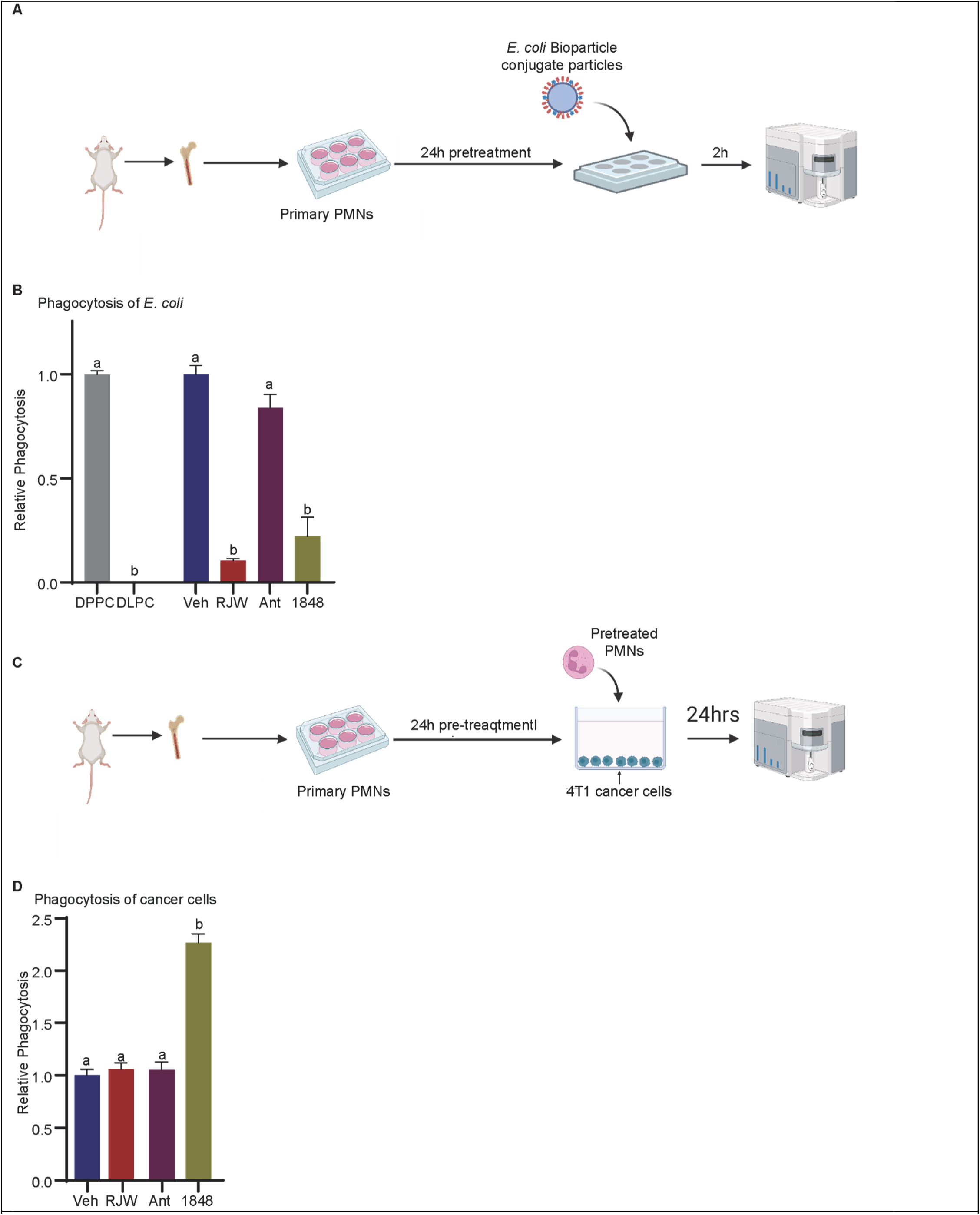
LRH-1 modulators regulate neutrophil phagocytosis. (**A**) Schematic overview of experimental setup for B. Neutrophils were treated with indicated ligands and evaluated for their ability to phagocytose *E. coli* conjugated to bioparticles. (**B**) Quantified data of neutrophils phagocytosing *E. coli* coated beads. Data was quantified and represented relative to respective controls. DLPC was normalized to control DPPC, and RR-RJW100 (RJW), LRH-1 antagonist (Ant) and the inverse agonist SR1858 (1848) were normalized to DMSO vehicle control (Veh). (**C**) Schematic overview of experimental setup for D. Neutrophils were treated with indicated ligands and evaluated for their ability to phagocytose 4T1 cancer cells. (**D**) Quantified data of neutrophils phagocytosing 4T1 cancer cells. Different letters denote statistically significant differences (P<0.05). DPPC vs DLPC was assessed with a two tailed student’s T test. The remaining treatment groups were assessed by one-way ANOVA followed by multiple comparison test with Šidáks correction. Schematics adapted from BioRender.

The canonical phagocytosis target for neutrophils is foreign pathogens such as *E. Coli*. However, neutrophils have been demonstrated to directly phagocytose cancer cells (55). To evaluate this, we cultured neutrophils that had been pre-treated with LRH-1 ligands with 4T1 mammary cancer cells, for 24h. The 4T1 cells had been stained with PKH26. After co-culture, neutrophils were then isolated and stained for Ly6G to confirm their cell-type and assessed for PKH26 positivity (outlined in **Fig. 5A**). Interestingly, RR-RJW100 or LRH-1 antagonist did not have a significant effect on phagocytosis of cancer cells at this timepoint (**Fig. 5C**). However, SR1848 increased phagocytosis of cancer cells (**Fig. 5C**). The effects of RR-RJW100 and SR1848 on phagocytosis of cancer cells was in stark contrast to those on phagocytosis of *E. coli* coated beads. This suggests that the effects of LRH-1 modulation are specific to the phagocytic target. Future work will be required to delineate the molecular mechanisms of these important observations.

### LRH-1 activity inhibits NETosis

NETosis and NETs have been implicated in cancer progression and reemergence from dormancy (56–59). Thus, it was important to evaluate whether LRH-1 activity regulated this aspect of neutrophil physiology. Initial analysis by image-capture flow cytometry (ImageStream) staining for extracellular DNA suggested that SR1848 increased NETosis, after priming with PMA (**Fig. 6A-B**). The phosphatidyl choline agonist DLPC decreased NETosis compared to its control DPPC, in neutrophils induced with PMA (**Fig. 6C**). To explore this relationship in more detail, we performed dose response curves for the LRH-1 antagonist, SR1848 and RR-RJW100. The LRH-1 agonist RR-RJW100 robustly inhibited NETosis in a dose related manner (**Fig. 6D**). On the other hand, both the antagonist and inverse agonist (SR1848) increased PMA-induced NETosis, in a dose dependent manner (**Fig. 6E-F**).

**Figure 6.**
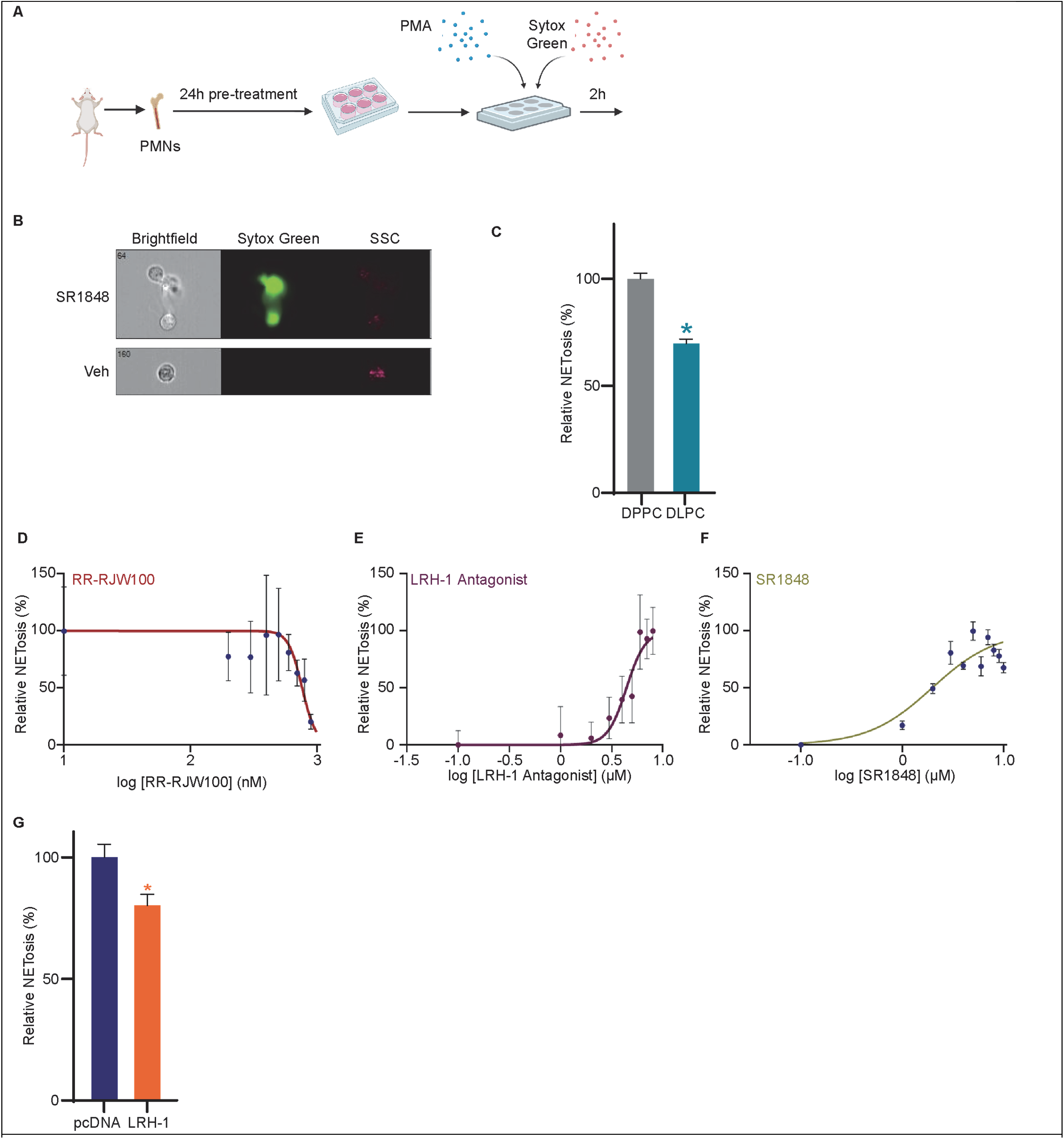
LRH-1 activity reduces neutrophil NETosis. (**A**) Experimental overview. Bone marrow derived neutrophils were harvested and cultured in the presence of ligands for 24h prior to induction of NETosis by PMA. NETosis was assessed by flow cytometry where extruded DNA was stained with SYTOX Green. (**B**) Image capture flow cytometry (ImageStream) confirmed NETosis post-PMA induction and that SR1848 increased it. (**C**) LRH-1 agonist DLPC reduces NETosis compared to its control DPPC (unpaired T test, P<0.05). (**D-F**) NETosis dose response curves for RR-RJW100, LRH-1 antagonist and SR1848 respectively. Replicate values at each concentration were averaged and then normalized from 0-100%. The normalized values were fit to a four-point non-linear regression model. (**G**) Neutrophils that were electroporated with an LRH-1 expression plasmid had decreased NETosis when induced with PMA (unpaired T test, P<0.05). 1µg of pcDNA control or LRH-1 expression plasmid were nucleofected. Schematics adapted from BioRender.

In addition to DNA, neutrophils also extrude associated proteins myeloiperoxidase (MPO) and a citrullinated histone H3 (CitH3). When we measured double-positive events for these proteins, we observed that the LRH-1 antagonist and SR1848 also increased these, confirming our conclusion that LRH-1 reduces the ability of neutrophils to undergo NETosis (**Supplementary Fig. 4A**). In order to substantiate these results, we used electroporation to successfully overexpress LRH-1 while maintaining viability (Nucleofector; mRNA quantification depicted in **Supplementary Fig. 4B**). In line with the impact of RR-RJW100, overexpression of LRH-1 in neutrophils reduced NETosis by approximately 20% relative to cells transfected with control pcDNA plasmid (**Fig. 6G**).

### LRH-1 in neutrophils supports CD4+ T cell expansion

Neutrophils and granulocytic myeloid derived suppressor cells have been implicated in the suppression of T cell expansion and activity (31,60,61). We observed that the LRH-1 antagonist or inverse agonist-primed neutrophils exhibited decreased surface expression of MHCII suggesting suppressed antigen-presenting capability (**Supplementary Fig. 5A**) (62–65). To investigate if LRH-1 activity in neutrophils could influence how they support T cells, we first made use of OTI and OTII mice, whose T cells have been engineered to recognize different ovalbumin (OVA) peptide antigens in the context of major histocompatibility class I and II molecules (H2-Kb or I-Ab) respectively, as we have used previously (14,17). Bone marrow derived neutrophils were treated with ligands for 24hrs prior to co-culture with OVA and OTI or OTII T cells (**Fig. 7A**). Pan-T cells from OTII mice demonstrated increased expansion when co-cultured with RR-RJW100 pretreated neutrophils compared to vehicle treated neutrophils, especially at later generations (**Fig. 7B**). This expansion was predominantly due to CD4+ cells as no effects were observed for CD8+ cells (**Fig. 7C** and **Supplementary Fig. 5B**). Further, T cells from OTI mice (OT1 being generally expressed in CD8+ cells) cells did not show any significant changes in subsequent expansion (**Supplementary Fig. 5C-E**). Pre-treatment with the LRH-1 antagonist modestly reduced T cell expansion, while the inverse agonist had no significant effects (**Fig. 7B-C**). Therefore, LRH-1 activity in neutrophils supports T cell expansion, especially helper T cell expansion (CD4+), when presented antigen from those neutrophils.

**Figure 7.**
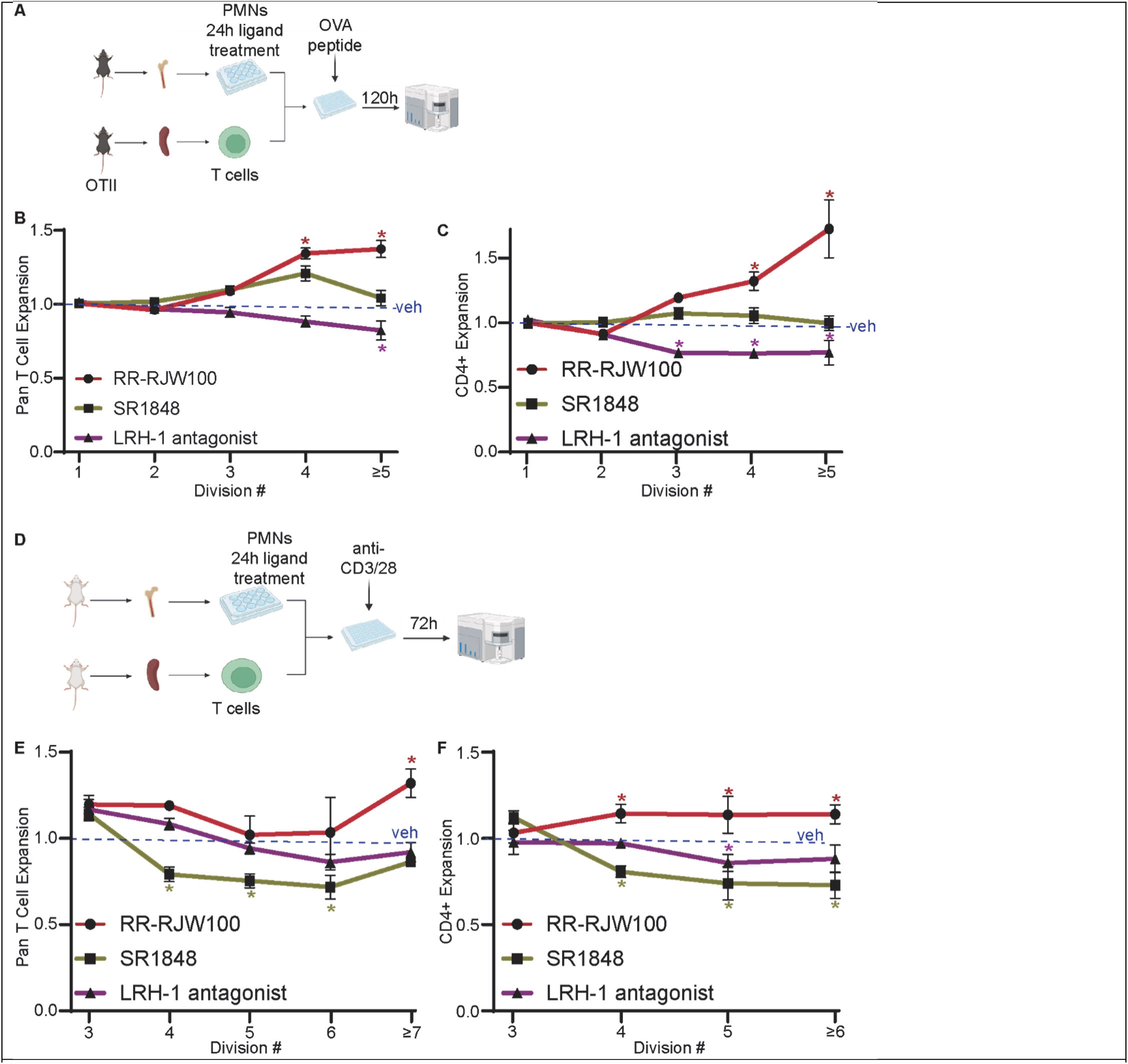
LRH-1 activity in neutrophils alters the expansion of co-cultured T cells. (**A**) Schematic overview of experimental setup for B&C. Neutrophils were isolated from the bone marrow of wildtype mice and cultured with indicated ligands, LPS (1mg/mL) prior to wash and co-culture with OVA peptide and T cells isolated from OTII mice. (**B**) Pan T cell expansion of OTII T cells was assessed by dilution of CFSE stain. Results are depicted as a ratio of expansion of treatment group to control (where vehicle control is normalized to 1 for each generation). (**C**) CD4+ T cell expansion of OTII T cells after co-culture with neutrophils treated as in (B). (**D**) Schematic overview of experimental setup for E&F. T cells from wildtype mice were activated with antibodies against CD3 and CD28 prior to their co-culture with pre-treated neutrophils. (**E**) Pan T cell expansion of pre-activated T cells after co-culture with neutrophils. (**F**) CD4+ T cell expansion of pre-activated T cells after co-culture with neutrophils, as in (F). Asterisks indicate statistically significant differences to vehicle control. Schematics adapted from BioRender.

However, these findings did not exclude the possibility that LRH-1 could also modulate neutrophil-T cell interactions through antigen presentation-independent mechanisms. To address this we co-cultured T cells that had been activated with antibodies against CD3 and CD28 prior to their co-culture with pre-treated neutrophils (**Fig. 7D**). In this experimental setup, RR-RJW100 continued to enhance T cell expansion at later divisions (**Fig. 7E**). Similarly, the increased expansion observed was largely restricted to CD4+ T cells (**Fig. 7F**). The LRH-1 antagonist and inverse agonist (SR1848) decreased expansion in this antigen-free experiment (**Fig. 7E-F**, and **Supplementary Fig. 5F**). Thus, LRH-1 activity in neutrophils serves to enhance T cell expansion, particularly CD4+ helper T cells. While these effects are evident in antigen-dependent contexts, they also persist when antigen presentation is bypassed, suggesting that LRH-1 may influence neutrophil T cell interactions through additional mechanisms.

Collectively, these data from *in vitro* / *ex vivo* systems suggest that LRH-1 activity shifts neutrophils towards a phenotype that is anti-cancer: maintaining phagocytosis, inhibiting NETosis and supporting CD4+ T cell expansion. Given our observations that the LRH-1 inverse agonist may decrease cancer cell proliferation, we next tested what effects were more important in terms of tumor growth: cancer cell intrinsic-versus neutrophil-mediated effects.

### LRH-1 agonist reduces ovarian tumor burden while an inverse agonist promotes mammary tumor growth

Our *in vitro* and *ex vivo* results strongly suggested that LRH-1 activity shifts neutrophils into an anti-tumor phenotype. However, the inverse agonist directly inhibited the proliferation of cancer cells themselves. Thus, it was important to evaluate the collective activities of LRH-1, in physiologically relevant *in vivo* models. Since DLPC is considered an endogenous LRH-1 agonist, we started by evaluating the effects of this compound. The pharmacokinetics of phosphatidylcholine structures such as DLPC are not well understood. Therefore, we tested this compound on a model of ovarian cancer, where DLPC could be administered by intraperitoneal injection, ensuring its tumoral exposure. Murine ID8 ovarian cancer cells bioengineered to express luciferase were grafted intraperitoneally into syngeneic C57BL/6 female mice and allowed to establish lesions for 7 days, at which point daily treatment with control-DPPC or DLPC was initiated (50mg/kg). Bioluminescent imaging revealed significant attenuation of growth in the peritoneal region with time (**Fig. 8A**). Interestingly DLPC also decreased metastatic burden in the lungs (**Supplementary Fig. 6**). Thus, regardless of its cancer cell intrinsic effects, LRH-1 activity reduces tumor burden.

**Figure 8.**
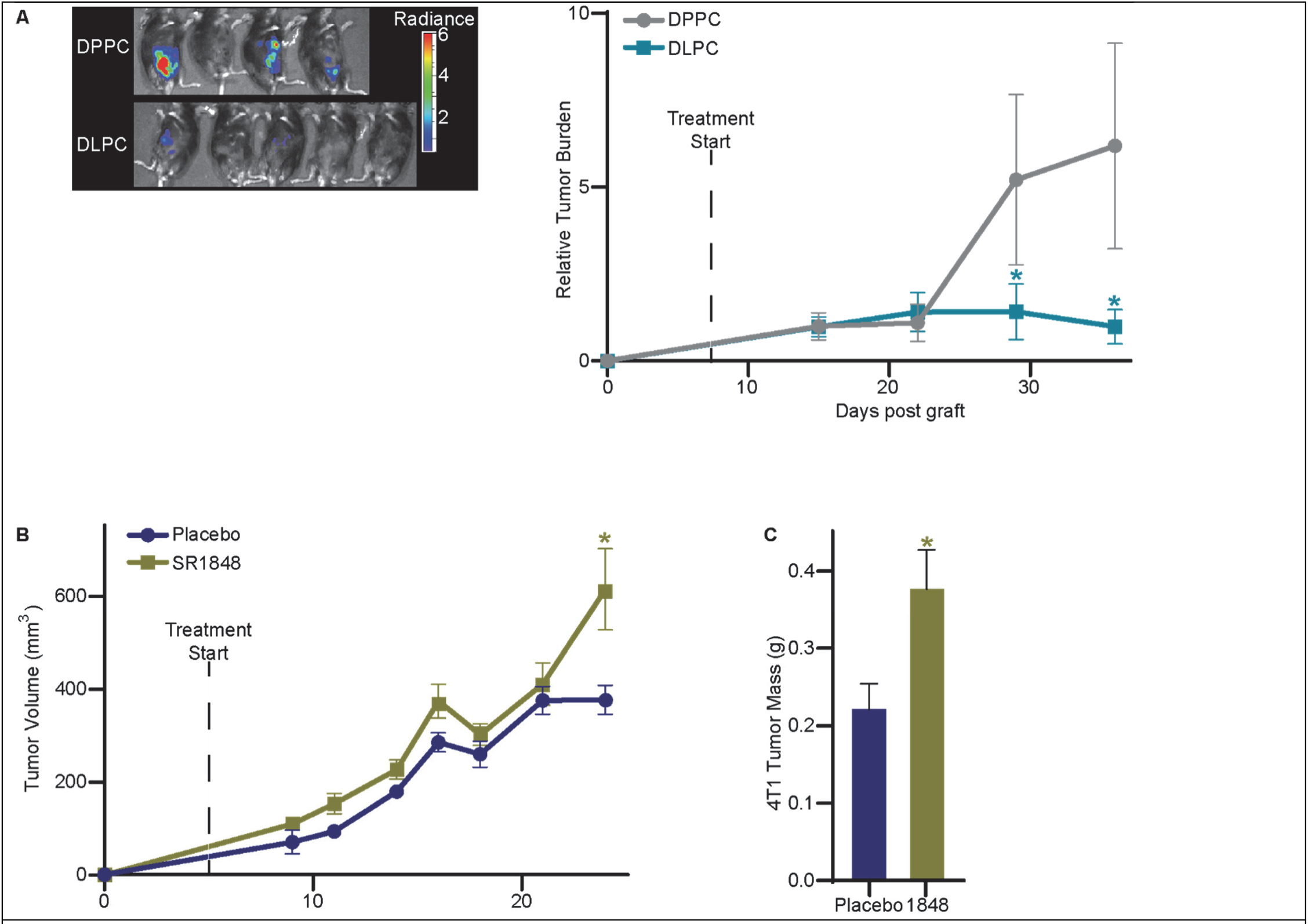
LRH-1 agonist reduces ovarian tumor growth while inverse agonist increases mammary tumor growth. (**A**) Murine ID8-luc ovarian cancer cells were grafted intraperitoneally and tumors allowed to establish lesions. 7 days post-graft, mice were randomized and treated with either control DPPC or the LRH-1 agonist DLPC (50mg/kg/day, 5 days/week). Bioluminescence imaging quantified tumor burden. Representative images are shown to the left of quantified data for the peritoneal region (corresponding data for the lung region can be found in **Supplementary** Fig. 6). Mean ± SEM is shown. Asterisks (*) denote statistically significant difference between DPPC control and DLPC for the indicated timepoint (data fit to a mixed-effects model followed by a post-hoc test with the Šidák’s correction for multiple comparisons). (**B**) Murine 4T1 mammary cancer cells were grafted orthotopically and tumor growth followed through time by direct caliper measurement. Daily treatment with placebo or 30mg/kg SR1848 started on day 5. (**C**) Resulting 4T1 tumor weights as assessed at necropsy. Asterisk (*) denotes statistically significant difference of weights tumors from DPPC-compared to DLPC-treated mice (P<0.05, unpaired, two-tailed T test).

We next evaluated the effects of the inverse agonist SR1848 on the growth of murine 4T1 mammary tumors, keeping in mind that 4T1 cellular proliferation was inhibited by this compound *in vitro* (**Fig. 2B**). Intriguingly, SR1848 modestly increased tumor growth compared to placebo in this fast-growing tumor model (**Fig. 8B**). Final tumor mass at necropsy was significantly higher in SR1848 treated mice (**Fig. 8C**). Even more robust growth-promoting effects of SR1848 were observed in the E0771 murine mammary tumor model (**Fig. 9**, described in more detail below). Thus, the cancer cell extrinsic effects of SR1848 dominate over its direct anti-proliferative activities, reinforcing the conclusion that LRH-1 activity in the tumor microenvironment suppresses tumor growth.

**Figure 9.**
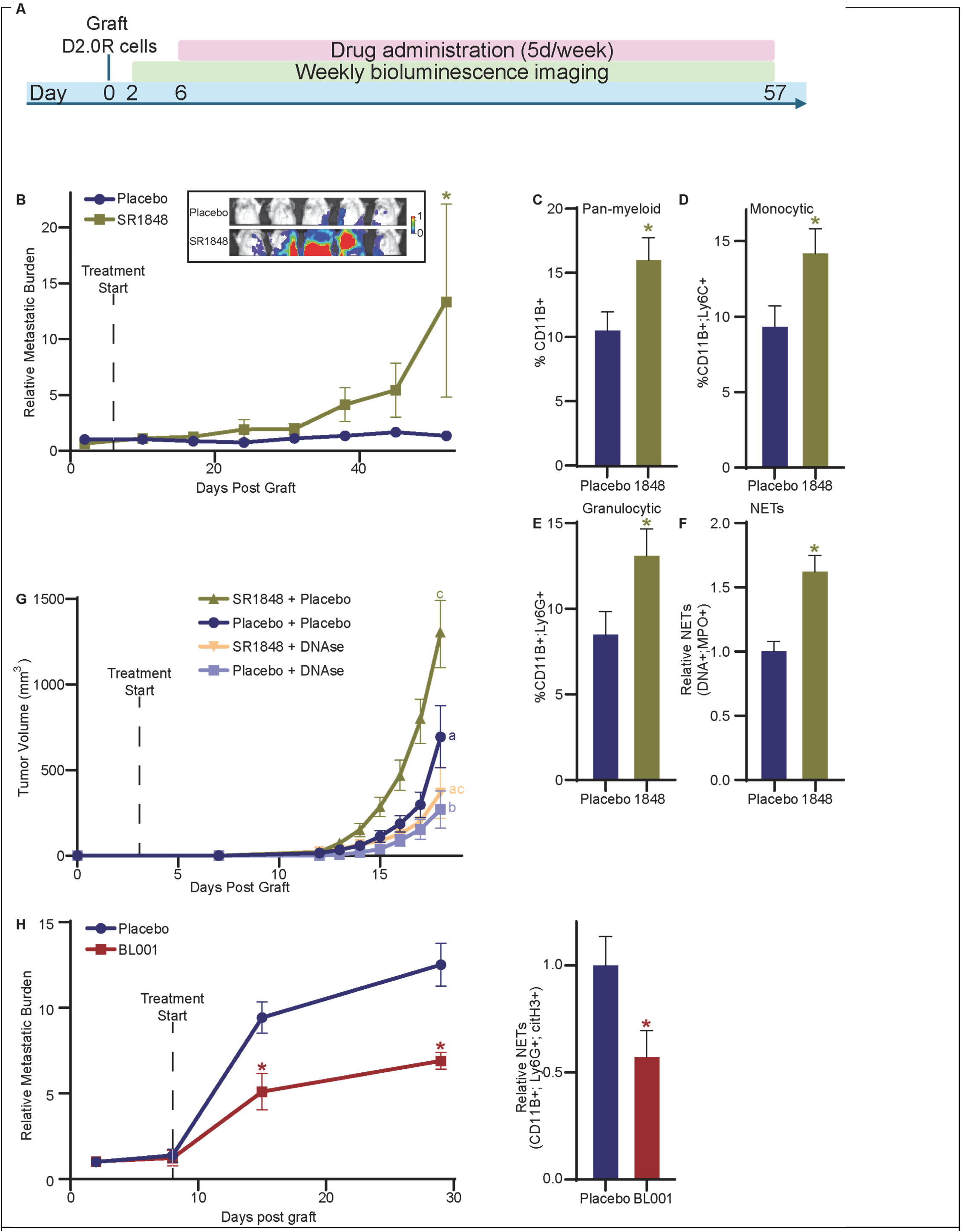
Inverse agonist of LRH-1 promotes recurrence from mammary cancer dormancy. (**A**) Schematic overview of experiment, the data of which is illustrated in B-F. Murine D2.0R cells were grafted intravenously and allowed to form dormant lesions. Six days post-graft, daily treatment with placebo or 30mg/kg SR1848 was initiated (5 days on, two days off). Recurrence was assessed by bioluminescent imaging. (**B**) Bioluminescent imaging through time. Placebo treated mice maintained dormant lesions while SR1848 treated mice experienced metastatic outgrowth. Representative images inset from quantified data. (**C-F**) Flow cytometric analysis of tumor myeloid immune cell infiltrate and NETs. (**C**) Pan myeloid immune cells (CD11b+). (**D**) Inflammatory monocytes or monocytic myeloid derived suppressor cells (CD11b+; Ly6C+). (**E**) Neutrophils, granulocytes or granulocytic myeloid derived suppressor cells (CD11b+; Ly6G+). (**F**) Relative presence of NETs as denoted by events that were positive for DNA and MPO. (**G**) SR1848 increases the growth of E0771 mammary tumors in a NET-dependent manner. Mice were orthotopically grafted with E0771 tumors. Two days post-graft, mice were treated daily with placebo SR1848 and/or DNAse I. Subsequent tumor outgrowth was measured. Different letters denote statistically significant differences (Two-Way ANOVA followed by multiple comparison test with Šidáks correction; differences only shown for the last time point). (**H**) LRH-1 agonist BL001 protects from reemergence from dormancy. D2.0R mammary cancer cells were grafted intravenously. 8 days post-graft mice were treated daily with placebo or 10mg/kg BL001 and 1mg/kg LPS to induce reemergence. Metastatic burden was quantified through time using bioluminescent imaging. Asterisks denote statistical significance (Two-Way ANOVA followed by multiple comparison test with Šidáks correction). (**I**) Relative presence of neutrophil-associated NETs as denoted by events that were positive for CD45, CD11b, Ly6G and citH3 (students T test).

### LRH-1 inhibits NETosis, protecting from mammary cancer reemergence from dormancy and reducing tumor growth

Metastasis involves a series of sequential steps including epithelial to mesenchymal transition, invasion and intravasation, cells in circulation, homing to specific tissues, extravasation, mesenchymal to epithelial transition, and subsequent outgrowth (66,67). Emerging clinical and experimental evidence suggests that the majority of breast cancer survivors continue to harbor cancer cells in distal tissues, that remain in a non-proliferative or dormant state (68–72). Through mechanisms that remain unclear, at some point dormant cells reemerge as growing metastatic lesions. It is metastatic disease that accounts for the vast majority of breast cancer related mortality.

Several recent reports have implicated neutrophils and NETosis in the reemergence from dormancy (56,57). Therefore, we evaluated the effects of SR1848 on a murine model of mammary cancer dormancy. D2.0R cells form dormant lesions in the lungs and bones when grafted intravenously. D2.0R cells were grafted and dormant lesions allowed to form (**Fig. 9A**). The lesions remained dormant in placebo treated mice. On the other hand, dormancy was broken in SR1848 treated mice (**Fig. 9B**). Flow cytometric analysis of SR1848-treated lungs revealed increased presence of myeloid cells in general (CD11B+; **Fig. 9C**), inflammatory monocytes (CD11B+;Ly6C+; **Fig. 9D**) and neutrophils (CD11B+;Ly6G+; **Fig. 9E**) cells, which is likely a reflection of increased tumor burden. Importantly however, was our observation of increased NETs within the lungs of SR1848-treated mice (as assessed by staining for extracellular DNA and MPO positive events; **Fig. 9F**). Thus, SR1848 increased NETosis and the likelihood of recurrence from dormancy.

Given previous reports implicating NETosis and tumor progression (73–75), it was necessary to evaluate the contributions of NETosis to the pro-tumor effects of SR1848. Thus, we investigated the effects of SR1848 on the tumor growth of E0771 mammary tumors, and assessed the requirement for NETs by administering DNAse I to digest any formed NETs. As expected, SR1848 robustly stimulated the growth of E0771 tumors in immune competent, syngeneic mice (**Fig. 9G**), similar to our findings in the 4T1 model (**Fig. 8B**). Interestingly, DNAse treatment alone attenuated tumor growth compared to placebo control and completely abrogated the stimulatory effects of SR1848 (**Fig. 9G**). This strongly suggests that the effects of SR1848 stimulate neutrophil NETosis resulting in tumor growth.

Because pharmacological LRH-1 activation reduced NETosis, preserved phagocytic capacity, and enhanced CD4+ T cell expansion in vitro, we next tested whether these immune-modulatory effects could be harnessed therapeutically *in vivo*. We therefore evaluated BL001, a clinically advanced LRH-1 agonist previously shown to be efficacious as an immunomodulator in murine models of Type 1 Diabetes (76,77). Given its potential use as a therapeutic, we evaluated whether BL001 could delay recurrence of mammary cancer. Mice were grafted with D2.0R cells and daily treatment started 8 days thereafter, along with a mild dose of LPS to trigger reemergence (57). In strong support of our hypothesis, BL001 treatment significantly reduced metastatic outgrowth (**Fig. 9G**). Consistent with this mechanism, lungs of BL001 treated mice exhibited reduced neutrophil-associated NETs in lungs of mice treated with BL001 (**Fig. 9I**). These findings position BL001 as a promising candidate to therapeutically exploit LRH-1 signaling for limiting metastatic recurrence.

## DISCUSSION

While immune checkpoint blockade (ICB) and CAR T cell therapies have demonstrated robust efficacy against certain cancer types, many remain refractory or quickly develop resistance to such approaches. For example, for metastatic breast cancer, ICB in combination with nab-paclitaxel is only approved for the treatment of TNBC tumors that also stain positive PD-L1. Even amongst this population, response rate is less than 25% (78,79). Resistance to ICB is multifactorial but a major contributor is thought to be from myeloid immune cells. Consequently, ‘re-educating’ myeloid cells is gaining traction as a therapeutic approach (18,80–83) . The complexity of myeloid cells has hampered efforts, with initial attempts at limiting myeloid cells or limiting their recruitment to the tumor microenvironment, ultimately failing (anti-CSF1 or CFSR) (83,84). This is likely due to the fact that myeloid cells are required to support a robust T cell response. Therefore, strategies to re-program myeloid cell functions rather than eliminate them, are likely to be more successful.

Our previous work has revealed that myeloid immune cells are particularly responsive to changes in cholesterol regulation (3). Many proteins involved in cholesterol homeostasis are nuclear receptors. This is important since nuclear receptors have a well-defined ligand binding domain and a demonstrated track-record of being targeted with small molecules; exemplified by the historic development of estrogen receptor modulators for the treatment of breast cancer (85–87).

LRH-1 is a nuclear receptor responsible for the regulation of cholesterol and lipid metabolism. We found that its mRNA expression within tumors is a favorable prognostic marker, being associated with an increase time to recurrence. However, the role of LRH-1 at the intersection of cancer biology and immune regulation in the tumor microenvironment remains underexplored. Many studies have found that LRH-1 activity in cancer cells promotes progression (reviewed in (3)). Indeed, data in this paper indicates that an LRH-1 antagonist or inverse agonist impaired the proliferation of different murine cancer cell lines. Yet, these observations were not congruent with the clinical data indicating that high LRH-1 expression in tumors was associated with increased survival time (**Figs. 1&2**). Thus, we speculated that cancer cell extrinsic effects of LRH-1 were likely predominant within the tumor microenvironment.

LRH-1 was also found to be relatively highly expressed in both human and mouse immune cells/cell lines particularly in granulocytes such as neutrophils. One caveat is that this evidence relies on mRNA expression data, and protein levels may differ. Neverthelss, neutrophils showed consistently high LRH-1 expression in healthy mammary tissue and breast tumors. Because neutrophils are increasingly recognized as central regulators of tumor progression, their high LRH-1 expression points to a potential role for this receptor in modulating pro- and/or anti-tumor activities.

The role of neutrophils in cancer biology is complex, imparting both pro- and anti-tumor programs (12,31). They are an abundant immune cell type within solid tumors, suggesting that strategies to manipulate their function may have translational potential. Using small molecule ligands for LRH-1, we found that LRH-1 activity regulates several neutrophil functions that are important for tumor biology: migration, phagocytosis, support of T cells and NETosis. For several functions, the regulation appears complex, whereby an agonist and inverse agonist had the same effect. This could be due to either (1) basal LRH-1 activity maintaining a function, and any perturbation in its basal activity resulting in decreased function, or (2) the selective modulation of LRH-1. Many nuclear receptors (if not all) can be selectively modulated – where ligands can have differing activities depending on the cellular context. This is best described for the estrogen and androgen receptors, but has also been described for the LXRs (15,87,88). In support of LRH-1 being selectively modulated are RNA-seq results post-ligand treatment, where there is significant DEG overlap between an agonist (RR-RJW100), antagonist and inverse agonist (SR1848). Furthermore, different ligand-binding result in unique exposed surfaces of LRH-1 and coregulator binding – the current paradigm for how nuclear receptor modulators exert their selective properties (89). Although the focus of future work, the nature of neutrophils makes it challenging to use functional genomics to test these two models of LRH-1 action.

There is evidence that neutrophils can directly phagocytose tumor cells, and also that tumor cells can escape from this process (55,90,91). Interestingly, both RR-RJW100 and SR1848 significantly reduced phagocytosis of *E. coli* bound beads. However, SR1848 increased phagocytosis of cancer cells. These apparently paradoxical findings suggest that the phagocytic programs evoked to consume *E. coli* vs. cancer cells are differentially regulated by LRH-1. These results are very intringing and warrant future investigation.

In addition to their well described roles in innate immunity, neutrophils have been suggested to exhibit regulatory effects on T cell expansion and activation in the case of different diseases (92–94). MPO-deficient neutrophils have been found to be correlated with the expansion of CD4+ cytotoxic T-lymphocytes (93). In addition, neutrophils have been demonstrated to suppress T cell proliferation and activation by acquiring inhibitory functions through either abnormal granulopoiesis, altering surface protein expression, or damaging the T cell membrane directly (92). Thus, we explored whether LRH-1 in neutrophils could regulate their interaction with T cells. Interestingly, T cells showed increased expansion when cocultured with neutrophils previously treated with an LRH-1 agonist, and reduced expansion in the presence of neutrophils pretreated with an LRH-1 antagonist or inverse agonist. This effect was particularly noticeable in CD4+ helper T cells during later divisions. The effects on T cell expansion were observed regardless of whether neutrophils activated the T cells through antigen presentation or when neutrophils were co-cultured with previously activated T cells. Thus, it is likely that a secreted or surface-expressed factor is responsible for the effects on T cells; something to be determined in future studies.

Our findings reveal a nuanced role for LRH-1 in regulating neutrophil–T cell interactions within the tumor microenvironment. *In vitro* treatment with LRH-1 antagonists or inverse agonists reduced MHC class II surface expression on neutrophils compared to LRH-1 agonist treatment, suggesting that LRH-1 activity enhances the antigen-presenting potential of these cells. Consistent with this, LRH-1 agonist–treated neutrophils promoted the expansion of CD4⁺ T cells in both CD3/CD28-activated pan-T cell cultures and antigen-specific OTII T cell co-cultures, highlighting LRH-1 as a positive regulator of neutrophil-mediated T cell activation. Interestingly, this immunostimulatory effect did not translate uniformly *in vivo*. In the 4T1 breast cancer model, treatment with an LRH-1 inverse agonist led to an increased frequency of Ly6G⁺ MHC II⁺ neutrophils in the lung, but no difference was observed in the primary tumor. These data suggest that the impact of LRH-1 modulation on neutrophil phenotype and function may vary by tissue microenvironment, with distinct consequences for local versus metastatic immune landscapes. Further studies are warranted to dissect the mechanisms driving these compartment-specific responses and to determine whether LRH-1 modulation could be leveraged therapeutically to enhance anti-tumor immunity.

Thus, several neutrophil functions relevant to tumor biology were altered by LRH-1 ligands *in vitro*, including migration, phagocytosis, T cell support and NETosis. When all these activities are at play simultaneously, in *in vivo* tumor models, it was found that an LRH-1 agonist decreased ovarian tumor growth while the LRH-1 inverse agonist increased tumor growth. The growth-promoting effects of the inverse agonist were, at least in part, mediated by NETs since DNAse treatment attenuated its effects. These findings are consistent with previous reports implicating NETs in cancer progression and reactivation of dormant cancer cells (56,57,95,96). Seminal studies have shown that NETs are involved in ‘sparking’ recurrence from dormancy (56–59). An LRH-1 inverse agonist reduced the time to recurrence in a murine model of mammary cancer dormancy. The dormant period between treatment of a primary tumor and recurrence offers a therapeutic window for preventative strategies, especially for breast cancer, where this period can last from months to decades. Importantly, the LRH-1 agonist BL001 was found to decrease recurrence and metastatic outgrowth in the D2.0R model of dormancy. BL001 had previously been shown to decrease hyperglycemia and immune-dependent inflammation in murine models of type 1 Diabetes Mellitus. It also decreased beta cell apoptosis in islets of type 2 diabetic patients (76). LRH-1 therefore exerts multifaceted effects on immunity, inflammation and tissue homeostasis, acting in neutrophils as well as other immune and non-immune compartments. Unlike strategies that deplete or broadly suppress myeloid cells, targeting LRH-1 offers a more selective means to reprogram neutrophil activity while preserving essential host-defense functions. Taken together, these findings position LRH-1 as a key regulator of neutrophil-mediated tumor control, with direct implications for metastasis and recurrence. The contrast between the protumor effects of LRH-1 inhibition and the protective role of BL001 underscores the therapeutic potential of selectively tuning this pathway rather than broadly suppressing neutrophil activity.

Future work will focus on the development of this axis for the prevention of recurrence. The ability to selectively modulate LRH-1 with small molecule ligands offers the opportunity to enhance neutrophil functions that limit tumor progression while preserving their essential roles in host defense and maintaining cholesterol and lipid metabolism. Such an approach could provide a novel strategy for the treatment of metastatic breast and ovarian cancers, as well as other solid tumors where neutrophil activity contributes to disease course. Strengthening innate immune responses to tumor cells and delaying or preventing metastatic relapse would represent a clinically meaningful advance; the present findings identifying LRH-1 in neutrophils as a target with tangible translational potential.

## MATERIALS AND METHODS

### Reagents

DNaseI was purchased from Sigma-Aldrich (catalogue number: D5025). 16:0 PC (DPPC) and 12:0 PC (DLPC) were purchased from Avanti Polar Lipids (850355 and 850335). SR1848 and RR-RJW100 were from MedChemExpress (HY-115613 and HY-131445B). LRH-1 antagonist from Sigma-Aldrich (5056010001). BL001 [(3a*S*,6a*R*)-1,2,3,3a,6,6a-hexahydro-4-(3-methoxyphenyl)-5-((*E*)-oct4-en-4-yl)-*N*-phenylpentalen-3a-amine] was synthesized by Sreeni Labs Private Limited (India), at a HPLC purity >98%. The semi-solid compound was dissolved in 100% DMSO at 20 μg/ml stock concentrations and further dissolved in WellSolve and used as previously described (76).

### Cell lines

4T1-luc and E0771 cells were a gift from Mark Dewhirst (Duke University School of Medicine). D2.0R cells were a gift from Michael Wendt (Iowa College of Medicine). ID8-luc cells were a gift from Robin Bachelder (Duke University School of Medicine). Cells were routinely tested for mycoplasma (Tumor Engineering and Phenotyping Shared Resource, Cancer Center at Illinois). We did not culture the cell line longer than 3 months after thawing the stocks.

### Animals

All procedure involving mice were approved by the Institutional Animal Care and Use Committee (IACUC) at the University of Illinois Urbana-Champaign. Wildtype C57BL/6 and BALB/C mice were purchased from Charles River Laboratory or Jackson Laboratories. Founder OTI and OTII mice were purchased from Jackson Laboratories and bred in-house in an institutional germ-free facility. ID8 tumor cells were grafted intraperitoneally. 4T1 and E0771 cells were grafted into the mammary fat pad. D2.0R cells were grafted intravenously. Bioluminescence acquisition was performed on a IVIS Spectrum CT live-animal imaging system as previously described (18).

### Preparation of primary polymorphonuclear neutrophil from bone marrow

Bone marrow cells were collected from mouse tibia and femur as described previously (97), passed through 70-μm cell strainer and red blood cells lysed in ACK buffer. Neutrophils were enriched using Ly6G beads (Miltenyi Biotec, 130120337) according to the manufacturer’s instructions.

### RNA-seq and bioinformatic analyses

#### Sample preparation and RNA isolation

Bone-marrow derived neutrophils were collected from 6 biological replicates of wildtype Balb/c per group. Total RNA was extracted using Quick-RNA Miniprep Plus Kit from Zymo Research following the manufacturer’s instructions. RNA integrity was assessed with an AATI Fragment. Analyzer, and samples with RNA quality number (RQN) ≥ 7 were used for library preparation.

#### Library preparation and sequencing

RNA-seq libraries were prepared using the Kapa Hyper Stranded mRNA library kit (Roche) with poly(A) selection. Libraries were sequenced on one 10B lane for 101 cycles from one end of the fragments on a NovaSeq X Plus with V1.0 sequencing kits.

#### Read alignment and quantification

Trimmed raw sequencing reads were quality-checked using MultiQC (v 1.28). Cleaned reads were aligned to the mouse reference genome GRCm39 and Annotation Release GCF_000001635.27-RS_2024_02 using STAR (v 2.7.10a) with default parameters. Gene-level counts were generated using featureCounts (Subread v 2.0.4).

#### Normalization and differential expression

<u>(A) DMSO vs. RR-RJW100 vs. LRH-1 antagonist</u>. Count data were imported into *edgeR* (v4.6.2) for preprocessing. Lowly expressed genes were filtered by retaining genes with counts per million (CPM) ≥ 1 in at least 3 samples. Data were normalized using the TMM method. Log-transformed CPM values were analyzed using *limma-trend*, with 1 RUV correction applied to account for unwanted variation. Differential expression was assessed using empirical Bayes moderated statistics, and p-values were adjusted for multiple testing using the Benjamini-Hochberg method. Genes with a false discovery rate (FDR) < 0.1 were considered significant. <u>(B) DMSO vs. SR1848.</u> Count data were imported into *edgeR* (v4.6.2) for downstream analysis. Lowly expressed genes were filtered by retaining those with counts per million (CPM) ≥ 0.5 in at least 3 samples. Data were normalized using the trimmed mean of M-values (TMM) method. Log-transformed CPM values were analyzed using *limma-trend*, and differential expression was assessed using the *treat* function to test for genes with a minimum absolute fold change threshold of 2. P-values were adjusted for multiple testing using the Benjamini-Hochberg method to control the false discovery rate (FDR). Genes with FDR < 0.005 were considered significant.

#### Pathway and functional analysis

Significantly differentially expressed genes were analyzed for pathway enrichment using *clusterProfiler* (v4.16.0) with KEGG and Gene Ontology (GO) databases. Visualization was performed using R packages including *ggplot2* (v3.5.2) and *gplots* (v3.2.0).

### Neutrophil-T cell co-culture assay

#### Wild type T cell -- CD3/CD28 activation

Neutrophils were seeded in a 96-well round bottom plate followed by treatment with DMSO, 1uM RR-RJW100, 5uM SR1848 or 10uM LRH-1 antagonist for 24hrs. CD3+ T cells were enriched from the spleen of wildtype BALB/c mice using the Dynabeads Untouched mouse T cells kit (Invitrogen, 11413D) according to the manufacturer’s instructions. T cells were labeled with the vital dye CFSE (BioLegend, 423801). T cells were co-cultured with neutrophils at a ratio of 1T:5N with 1mg/mL CD3 (BD Pharmingen™, 553057) and 1mg/mL CD28 antibody (BD Pharmingen™, 553294), for 72hrs. The proliferation of T cells (CFSE) and differentiation (CD4+/CD8+) were assessed using flow cytometry.

#### OTII T cell -- with OVA_323-339_

Neutrophils were prepared from C57BL/6 mice as described. T cells were co-cultured with washed neutrophils at a ratio of 5T:1N with 10ug/mL of OVA_323-229_ peptide (Bachem, 4034255), for 120hrs.

### NETosis

1. <u>Dose response assay –*Detection of extruded DNA using SYTOX green.*</u> Neutrophils were seeded at a density and treated for 24hrs. PMA, a NETosis inducer, was added at a final concentration of 30nM, along with diluted SYTOX Green stain (1:250 dilution, Invitrogen S7020). After 2hrs of incubation, percentage of events that stained positive for SYTOX Green were quantified using flow cytometry.
2. *<u>Stain for citrullinated histone 3 (cith3) and MPO after PMA stimulation</u>.* Treated neutrophils were stained with Ly6G (FITC), MPO (AF647) and citH3 (primary antibody – Abcam, ab281584; secondary antibody – Invitrogen, A-11011) and quantified for the percentage of positively stained cells using flow cytometry.

### Phagocytosis assay

After treatment neutrophils were cultured with pHrodo conjugated E.coli bioparticles (Invitrogen, P35360) for 2hrs. Phagocytosis activity was quantified according to the percentage of cells that had engulfed the conjugated bioparticles using flow cytometry.

### Migration

Pre-treated neutrophils were washed and transferred to the transwell inserts. The cells were allowed to migrate towards cancer-cell conditioned media for 2hrs. Migrated cells were collected and stained for surface Ly6G and quantified using flow cytometry.

### Nucleofection

Neutrophils were nucleofected with 1ug of plasmid DNA (pcDNA and LRH-1 plasmid) using Human Monocyte Nucleofector® Kit (Lonza VPA-1007) on Amaxa Nucleofector 2D according to the manufacturer’s instructions.

### MHC II regulation

Neutrophils were stained for MHC II/IA-IE quantified for the percentage of positively stained cells using flow cytometry.

### Real-time quantitative polymerase chain reaction (RT-qPCR)

RNA was extracted using the GeneJet RNA Purification kit (Thermo Fisher) (Catalog no. K0731), reverse transcribed and quantified through qPCR as described (98). Relative expression was determined via the 2–ΔΔCT method and normalized to the TATA box binding protein (TBP), a housekeeping gene.

### Data analysis and statistics

All general statistical analyses were performed using GraphPad Prism software. Data were analyzed using two-sided t-tests, one-way or two-way ANOVA, followed by Dunnett’s, Šidák’s, Tukey or Bonferroni’s multiple comparisons test, as indicated in the figure legends. Statistical significance was determined as P < 0.05.

## Supporting information

Supplementary Material, Wang et al.

## Acknowledgments

We would like to thank the patients whose tumors populated data in the TCGA, METABRIC and Human Protein Atlas initiatives. We would also like to thank our breast cancer advocate team: Sarah Adams, Renaé Strawbridge, Jamie Holloway, Lea Ann Carson, Susan Stewart and Catherine Applegate. The Tumor Engineering and Phenotyping Core at the Cancer Center at Illinois provided mycoplasma testing. The Roy J. Carver Biotechnology Center performed the RNA-sequencing.

## Funding

Department of Defense BC241117/HT9425-25-1-0285 (ERN)

Department of Defense Era of Hope Scholar Award BC200206/W81XWH-20-BCRP-EOHS (ERN)

National Institutes of Health grant R01 CA288207 (ERN)

National Institutes of Health grant R01 CA234025 (ERN)

National Institutes of Health grant T32 GM136629 (HEVG)

National Institutes of Health grant T32 ES007326 (ATN)

National Institutes of Health grant T32 EB019944 (CPS)

Vencer el Cancer (BRG)

DiabetesCero Foundation (BRG)

Breakthrough T1D formerly Juvenile Diabetes Research Foundation (2-SRA-2019-837-S-B, 3-SRA-2023-1307-S-B to B.R.G)

The Endocrine Society Research Experiences for Graduate and Medical Students Award (SVB) The Cancer Center at Illinois Graduate Cancer Scholarship Program Award (SVB)

Postdoctoral Fellows Program at the Beckman Institute for Advanced Science and Technology (NK) University of Illinois (ERN)

## Data and materials availability

Data and/or reagents will be provided upon reasonable request. RNA-seq and ATAC-seq data will be uploaded to GEO upon acceptance of this manuscript.

