## Supplementary Material, Wang et al. for "LRH-1 is a novel regulator of neutrophil-driven immune responses within the tumor microenvironment"

**Affiliations:**
[1] Department of Molecular and Integrative Physiology, University of Illinois Urbana Champaign, Illinois, USA.
[2] Beckman Institute for Advanced Science and Technology, University of Illinois Urbana Champaign, Illinois, USA
[3] Roy J. Carver Biotechnology Center, University of Illinois Urbana Champaign, Urbana Illinois, 61801 USA.
[4] Andalusian Center of Molecular Biology and Regenerative Medicine-CABIMER, Junta de Andalucía-University of Pablo de Olavide-University of Seville-CSIC, Seville, Spain
[5] Division of Nutritional Sciences, University of Illinois Urbana Champaign, Illinois, USA
[6] Carl R. Woese Institute for Genomic Biology- Anticancer Discovery from Pets to People, University of Illinois Urbana Champaign, Illinois, USA
[7] Cancer Center at Illinois, University of Illinois Urbana Champaign, Illinois, USA

**Keywords:**
LRH-1, liver receptor homolog 1, NR5A2, nuclear receptor, cholesterol, myeloid cell, neutrophil, breast cancer, ovarian cancer

**Acknowledgments:**
We would like to thank the patients whose tumors populated data in the TCGA, METABRIC and Human Protein Atlas initiatives. We would also like to thank our breast cancer advocate team: Sarah Adams, Renaé Strawbridge, Jamie Holloway, Lea Ann Carson, Susan Stewart and Catherine Applegate. The Tumor Engineering and Phenotyping Core at the Cancer Center at Illinois provided mycoplasma testing. The Roy J. Carver Biotechnology Center performed the RNA-sequencing.

**Funding:**
Department of Defense BC241117/HT9425-25-1-0285 (ERN)
Department of Defense Era of Hope Scholar Award BC200206/W81XWH-20-BCRP-EOHS (ERN)
National Institutes of Health grant R01 CA288207 (ERN)
National Institutes of Health grant R01 CA234025 (ERN)
National Institutes of Health grant T32 GM136629 (HEVG)
National Institutes of Health grant T32 ES007326 (ATN)
National Institutes of Health grant T32 EB019944 (CPS)
Vencer el Cancer (BRG)
DiabetesCero Foundation (BRG)
Breakthrough T1D formerly Juvenile Diabetes Research Foundation (2-SRA-2019-837-S-B, 3-SRA-2023-1307-S-B to B.R.G)
The Endocrine Society Research Experiences for Graduate and Medical Students Award (SVB)
The Cancer Center at Illinois Graduate Cancer Scholarship Program Award (SVB)
Postdoctoral Fellows Program at the Beckman Institute for Advanced Science and Technology (NK)
University of Illinois (ERN)

**Data and materials availability:** Data and/or reagents will be provided upon reasonable request. RNA-seq and ATAC-seq data will be uploaded to GEO upon acceptance of this manuscript.

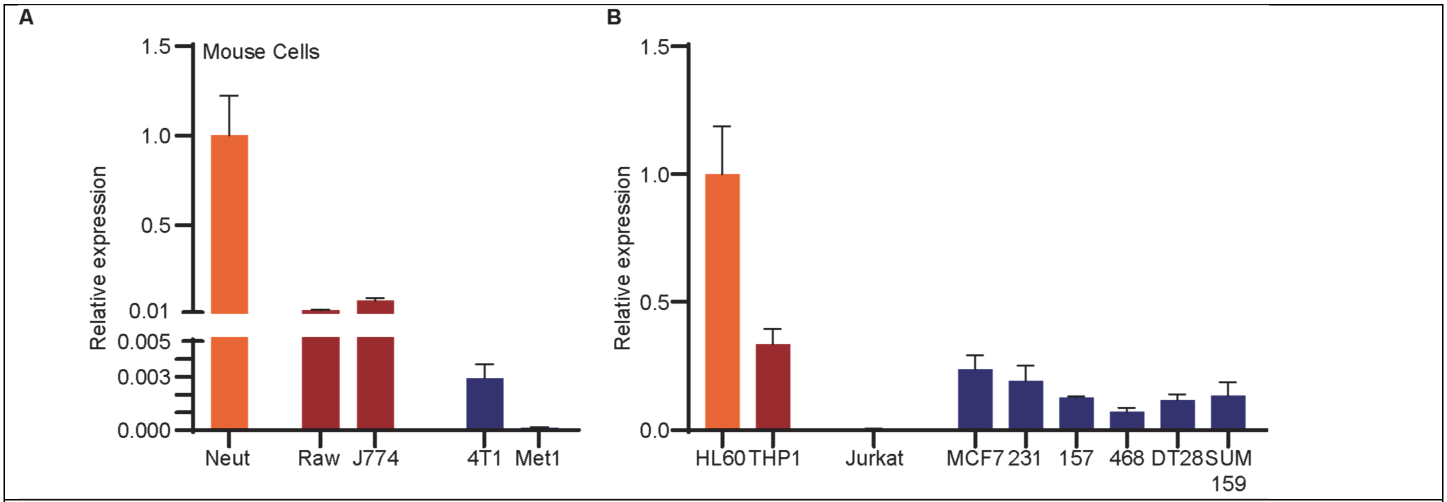


**Supplementary Figure 1.** ***LRH-1 is expressed at relatively high levels in neutrophils or neutrophil-like cell lines***. (**A**) Murine cells. Primary murine neutrophils (neut) were isolated from bone marrow of C57BL/6 female mice. RAW264.7 (RAW) and J774A.1 (J774) cells are monocytic/macrophage-like cell lines. 4T1 and Met1 are murine mammary cancer lines that model TNBC. RNA was isolated from cultured cells and LRH-1 mRNA assessed by qPCR. Note the split y-axis. (**B**) Human cells. HL60 are a leukemia line resembling neutrophils. THP1 cells are a monocytic leukemia cell line. Jurkat cells are immortalized T cells. MCF7 are ERα+ breast cancer cells. MDA MB 231, 157 and 468, and DT28 and SUM159 are TNBC cells.

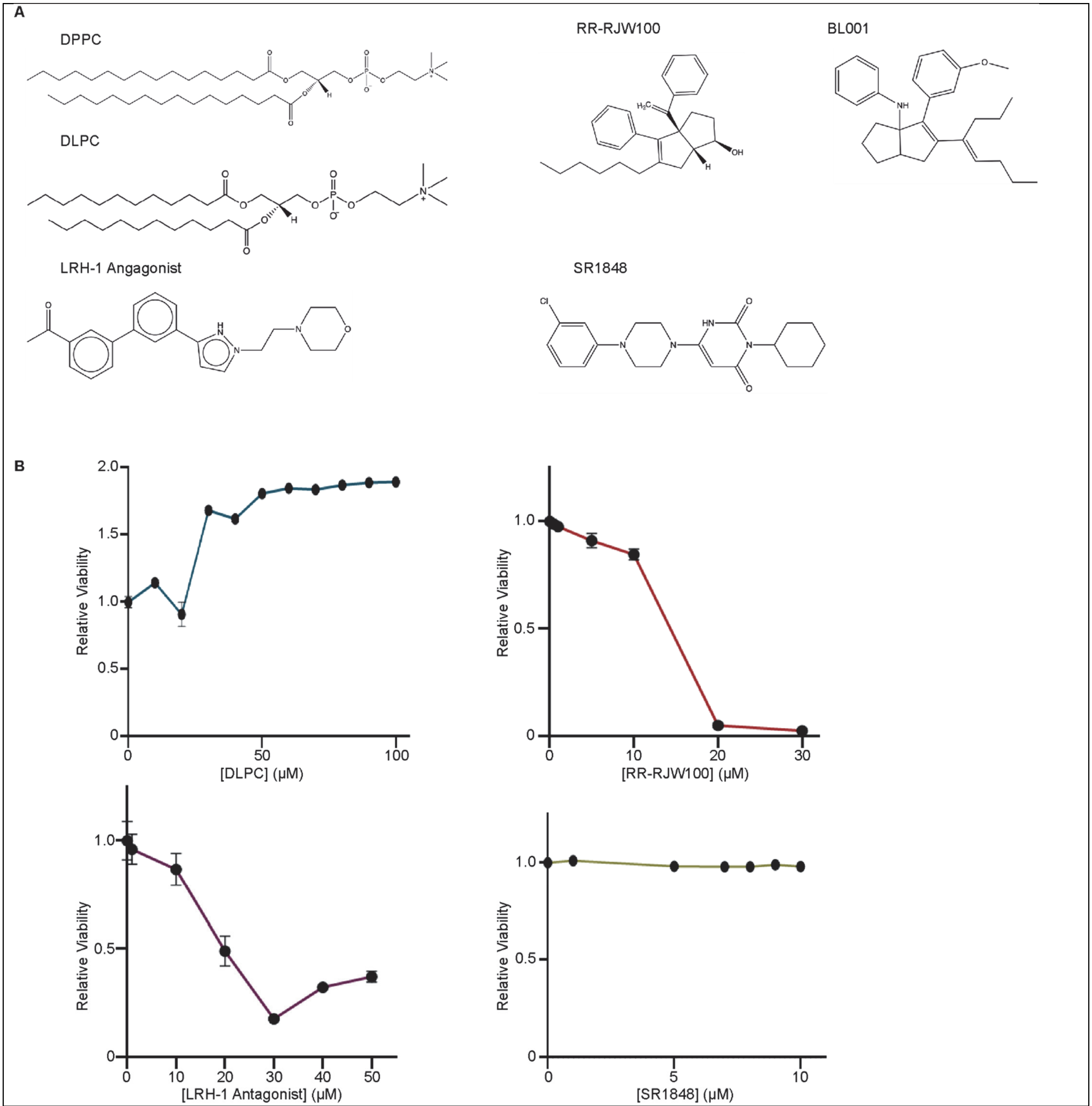


**Supplementary Figure 2.** ***Small molecule modulators of LRH-1 have relatively low cytotoxicity.*** (**A**) Chemical structures of the small molecule modulators of LRH-1 used in this study (modified from ChemDraw). (**B**) Primary PMNs were isolated, seeded at a density of 400k cell/well in 12-well non-treated plates and treated with the indicated ligands for 24hrs. Cells were then stained for PI and Annexin V according to the manufacturer's instructions. Relative viability was ascertained as live cells normalized to those in the vehicle (0µM group).

A

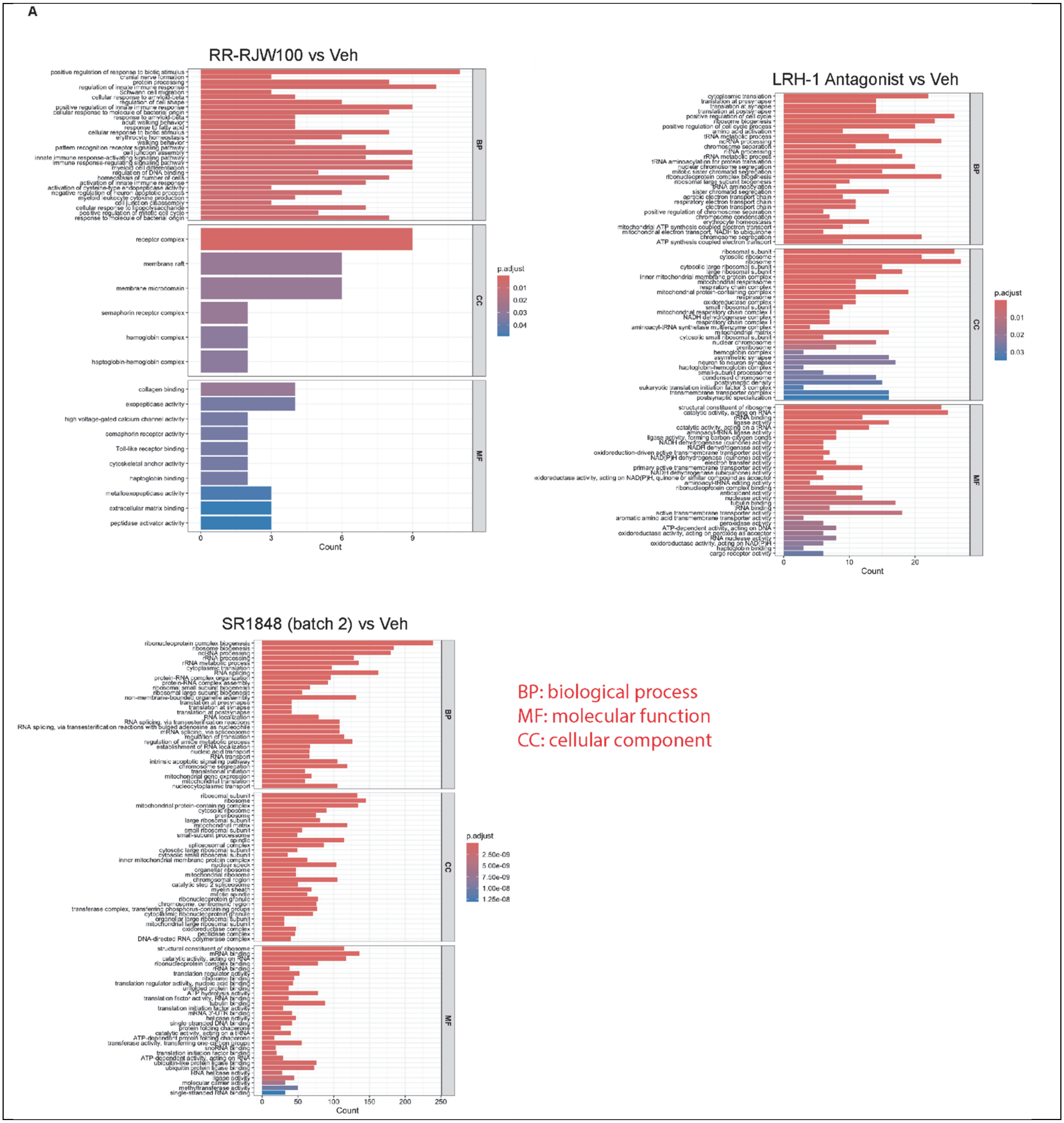
RR-RJW100 vs Veh
receptor complex
membrane raft
membrane microdomain
semaphorin receptor complex
hemoglobin complex
haptoglobin-hemoglobin complex
collagen binding
exopeptidase activity
high voltage-gated calcium channel activity
semaphorin receptor activity
Toll-like receptor binding
cytoskeletal anchor activity
haptoglobin binding
metalloexopeptidase activity
extracellular matrix binding
peptidase activator activity
Count
p.adjust
LRH-1 Antagonist vs Veh
Count
p.adjust
SR1848 (batch 2) vs Veh
Count
p.adjust
BP
CC
MF
BP: biological process
MF: molecular function
CC: cellular component

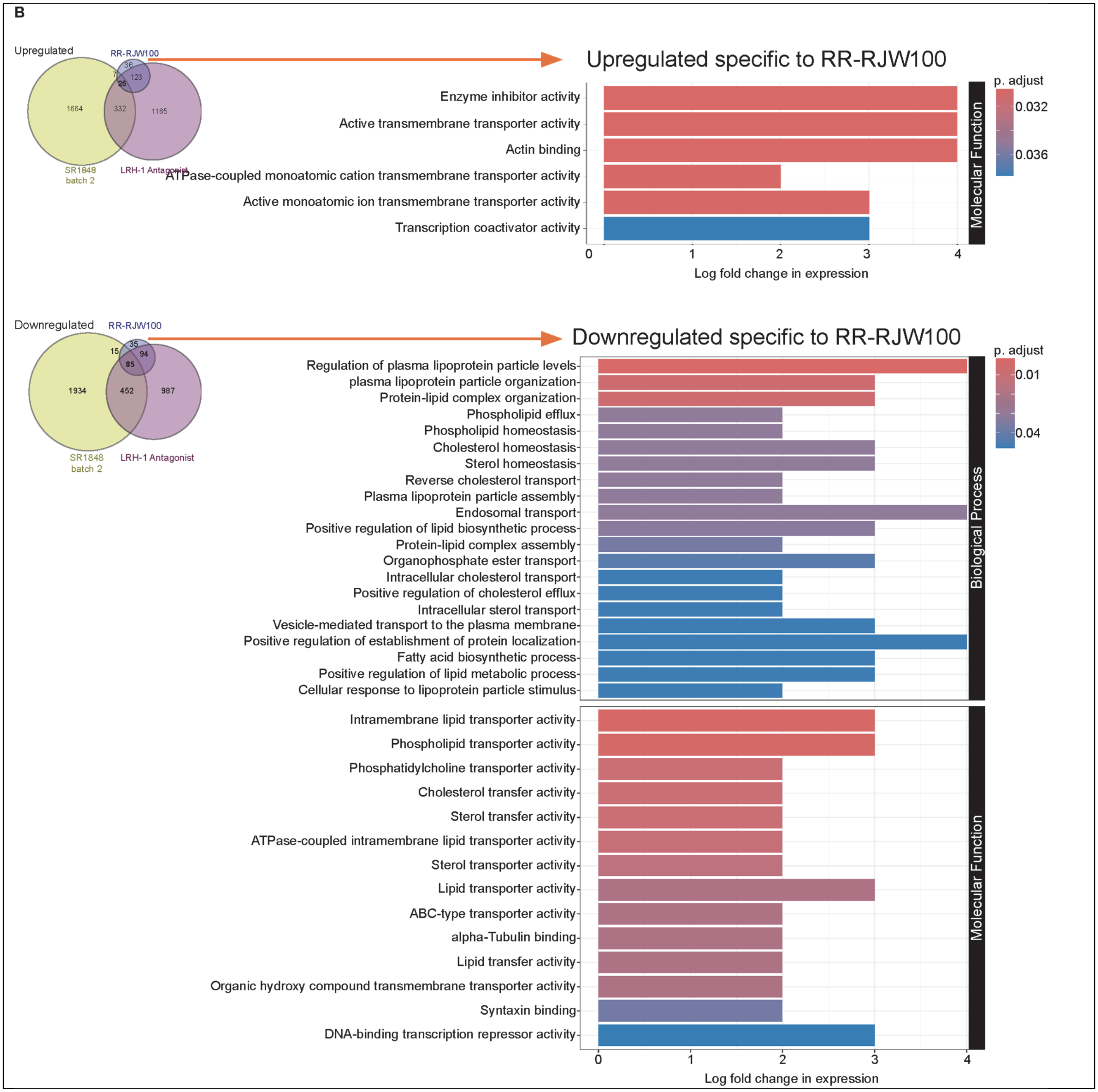

B
Upregulated
RR-RJW100
36
7
25
123
1664
332
1165
SR1848 batch 2
LRH-1 Antagonist
Upregulated specific to RR-RJW100
Enzyme inhibitor activity
Active transmembrane transporter activity
Actin binding
ATPase-coupled monoatomic cation transmembrane transporter activity
Active monoatomic ion transmembrane transporter activity
Transcription coactivator activity
Molecular Function
p. adjust
0.032
0.036
0 1 2 3 4
Log fold change in expression
Downregulated
RR-RJW100
35
15
94
85
1934
452
987
SR1848 batch 2
LRH-1 Antagonist
Downregulated specific to RR-RJW100
Regulation of plasma lipoprotein particle levels
plasma lipoprotein particle organization
Protein-lipid complex organization
Phospholipid efflux
Phospholipid homeostasis
Cholesterol homeostasis
Sterol homeostasis
Reverse cholesterol transport
Plasma lipoprotein particle assembly
Endosomal transport
Positive regulation of lipid biosynthetic process
Protein-lipid complex assembly
Organophosphate ester transport
Intracellular cholesterol transport
Positive regulation of cholesterol efflux
Intracellular sterol transport
Vesicle-mediated transport to the plasma membrane
Positive regulation of establishment of protein localization
Fatty acid biosynthetic process
Positive regulation of lipid metabolic process
Cellular response to lipoprotein particle stimulus
Biological Process
Intramembrane lipid transporter activity
Phospholipid transporter activity
Phosphatidylcholine transporter activity
Cholesterol transfer activity
Sterol transfer activity
ATPase-coupled intramembrane lipid transporter activity
Sterol transporter activity
Lipid transporter activity
ABC-type transporter activity
alpha-Tubulin binding
Lipid transfer activity
Organic hydroxy compound transmembrane transporter activity
Syntaxin binding
DNA-binding transcription repressor activity
Molecular Function
p. adjust
0.01
0.04
0 1 2 3 4
Log fold change in expression

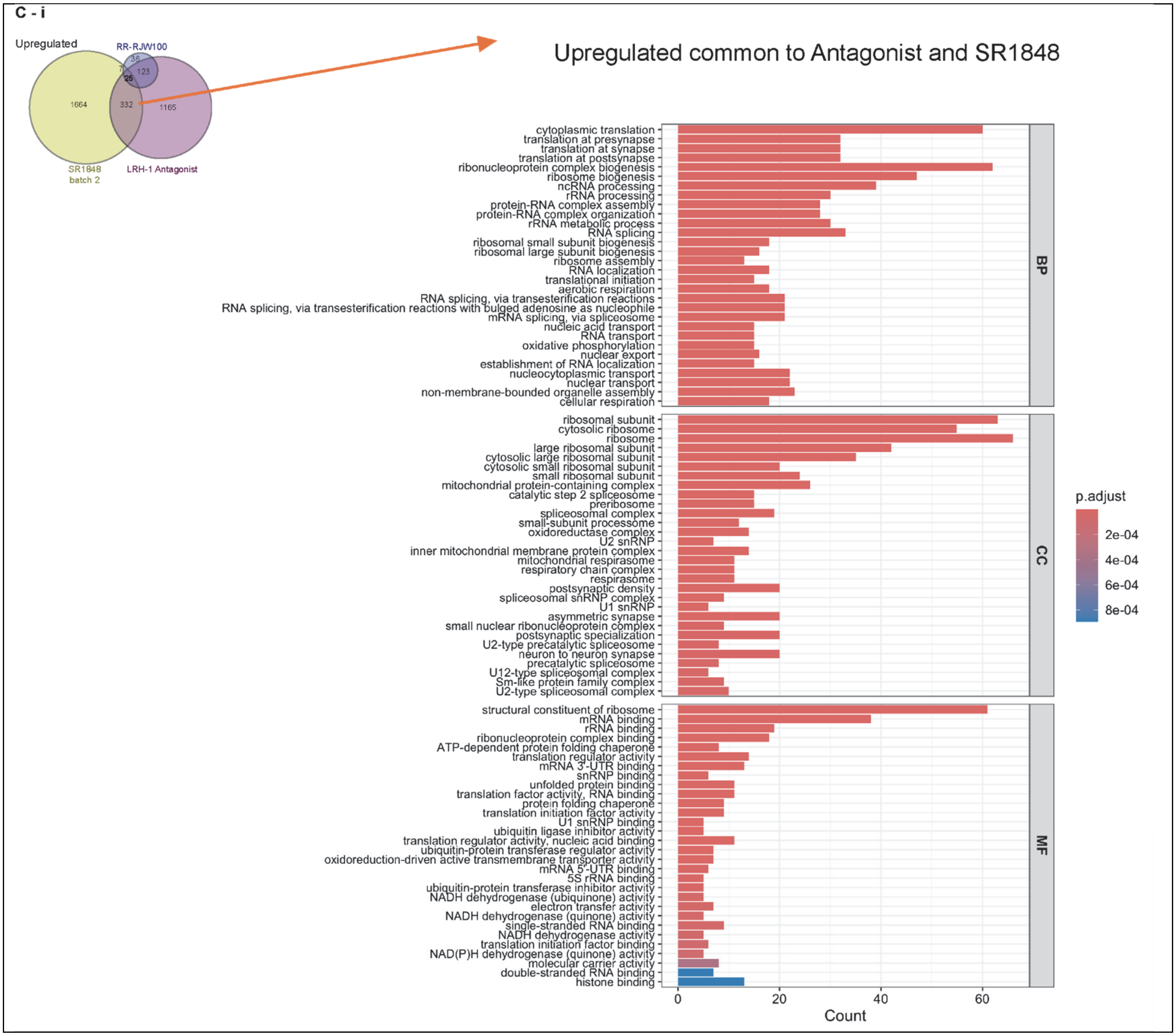
C - i
Upregulated
RR-RJW100
SR1848 batch 2
LRH-1 Antagonist
1664
332
1165
123
36
25
7
Upregulated common to Antagonist and SR1848
cytoplasmic translation
translation at presynapse
translation at synapse
translation at postsynapse
ribonucleoprotein complex biogenesis
ribosome biogenesis
ncRNA processing
rRNA processing
protein-RNA complex assembly
protein-RNA complex organization
rRNA metabolic process
RNA splicing
ribosomal small subunit biogenesis
ribosomal large subunit biogenesis
ribosome assembly
RNA localization
translational initiation
aerobic respiration
RNA splicing, via transesterification reactions
RNA splicing, via transesterification reactions with bulged adenosine as nucleophile
mRNA splicing, via spliceosome
nucleic acid transport
RNA transport
oxidative phosphorylation
nuclear export
establishment of RNA localization
nucleocytoplasmic transport
nuclear transport
non-membrane-bounded organelle assembly
cellular respiration
BP
ribosomal subunit
cytosolic ribosome
ribosome
large ribosomal subunit
cytosolic large ribosomal subunit
cytosolic small ribosomal subunit
small ribosomal subunit
mitochondrial protein-containing complex
catalytic step 2 spliceosome
preribosome
spliceosomal complex
small-subunit processome
oxidoreductase complex
U2 snRNP
inner mitochondrial membrane protein complex
mitochondrial respirasome
respiratory chain complex
respirasome
postsynaptic density
spliceosomal snRNP complex
U1 snRNP
asymmetric synapse
small nuclear ribonucleoprotein complex
postsynaptic specialization
U2-type precatalytic spliceosome
neuron to neuron synapse
precatalytic spliceosome
U12-type spliceosomal complex
Sm-like protein family complex
U2-type spliceosomal complex
CC
structural constituent of ribosome
mRNA binding
rRNA binding
ribonucleoprotein complex binding
ATP-dependent protein folding chaperone
translation regulator activity
mRNA 3'-UTR binding
snRNP binding
unfolded protein binding
translation factor activity, RNA binding
protein folding chaperone
translation initiation factor activity
U1 snRNP binding
ubiquitin ligase inhibitor activity
translation regulator activity, nucleic acid binding
ubiquitin-protein transferase regulator activity
oxidoreduction-driven active transmembrane transporter activity
mRNA 5'-UTR binding
5S rRNA binding
ubiquitin-protein transferase inhibitor activity
NADH dehydrogenase (ubiquinone) activity
electron transfer activity
NADH dehydrogenase (quinone) activity
single-stranded RNA binding
NADH dehydrogenase activity
translation initiation factor binding
NAD(P)H dehydrogenase (quinone) activity
molecular carrier activity
double-stranded RNA binding
histone binding
MF
0
20
40
60
Count
p.adjust
2e-04
4e-04
6e-04
8e-04

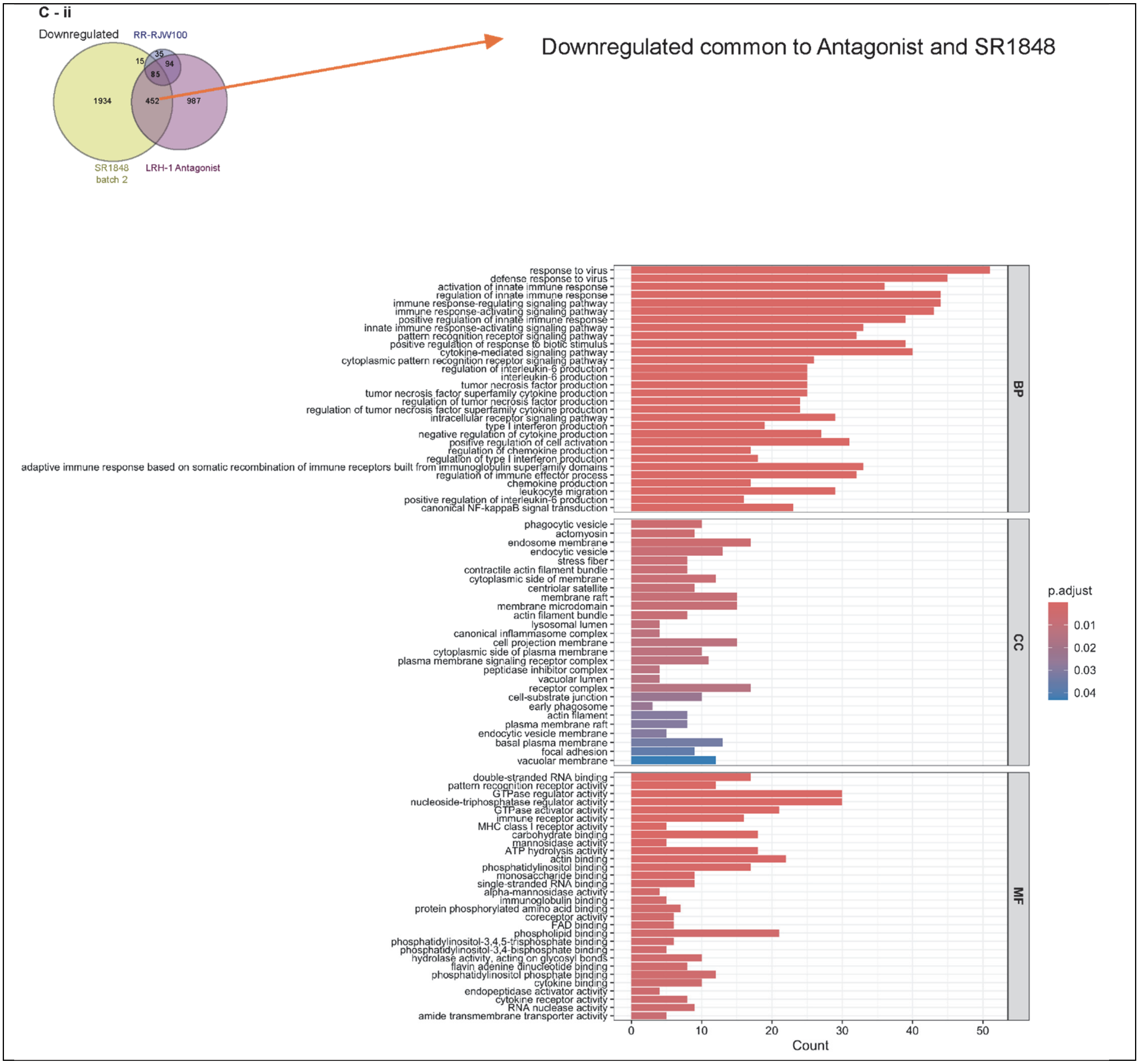

C - ii
Downregulated
RR-RJW100
15
35
94
85
1934
452
987
SR1848 batch 2
LRH-1 Antagonist
Downregulated common to Antagonist and SR1848
response to virus
defense response to virus
activation of innate immune response
regulation of innate immune response
immune response-regulating signaling pathway
immune response-activating signaling pathway
positive regulation of innate immune response
innate immune response-activating signaling pathway
pattern recognition receptor signaling pathway
positive regulation of response to biotic stimulus
cytokine-mediated signaling pathway
cytoplasmic pattern recognition receptor signaling pathway
regulation of interleukin-6 production
interleukin-6 production
tumor necrosis factor production
tumor necrosis factor superfamily cytokine production
regulation of tumor necrosis factor production
regulation of tumor necrosis factor superfamily cytokine production
intracellular receptor signaling pathway
type I interferon production
negative regulation of cytokine production
positive regulation of cell activation
regulation of chemokine production
regulation of type I interferon production
adaptive immune response based on somatic recombination of immune receptors built from immunoglobulin superfamily domains
regulation of immune effector process
chemokine production
leukocyte migration
positive regulation of interleukin-6 production
canonical NF-kappaB signal transduction
phagocytic vesicle
actomyosin
endosome membrane
endocytic vesicle
stress fiber
contractile actin filament bundle
cytoplasmic side of membrane
centriolar satellite
membrane raft
membrane microdomain
actin filament bundle
lysosomal lumen
canonical inflammasome complex
cell projection membrane
cytoplasmic side of plasma membrane
plasma membrane signaling receptor complex
peptidase inhibitor complex
vacuolar lumen
receptor complex
cell-substrate junction
early phagosome
actin filament
plasma membrane raft
endocytic vesicle membrane
basal plasma membrane
focal adhesion
vacuolar membrane
double-stranded RNA binding
pattern recognition receptor activity
GTPase regulator activity
nucleoside-triphosphatase regulator activity
GTPase activator activity
immune receptor activity
MHC class I receptor activity
carbohydrate binding
mannosidase activity
ATP hydrolysis activity
actin binding
phosphatidylinositol binding
monosaccharide binding
single-stranded RNA binding
alpha-mannosidase activity
immunoglobulin binding
protein phosphorylated amino acid binding
coreceptor activity
FAD binding
phospholipid binding
phosphatidylinositol-3,4,5-trisphosphate binding
phosphatidylinositol-3,4-bisphosphate binding
hydrolase activity, acting on glycosyl bonds
flavin adenine dinucleotide binding
phosphatidylinositol phosphate binding
cytokine binding
endopeptidase activator activity
cytokine receptor activity
RNA nuclease activity
amide transmembrane transporter activity
BP
CC
MF
0
10
20
30
40
50
Count
p.adjust
0.01
0.02
0.03
0.04

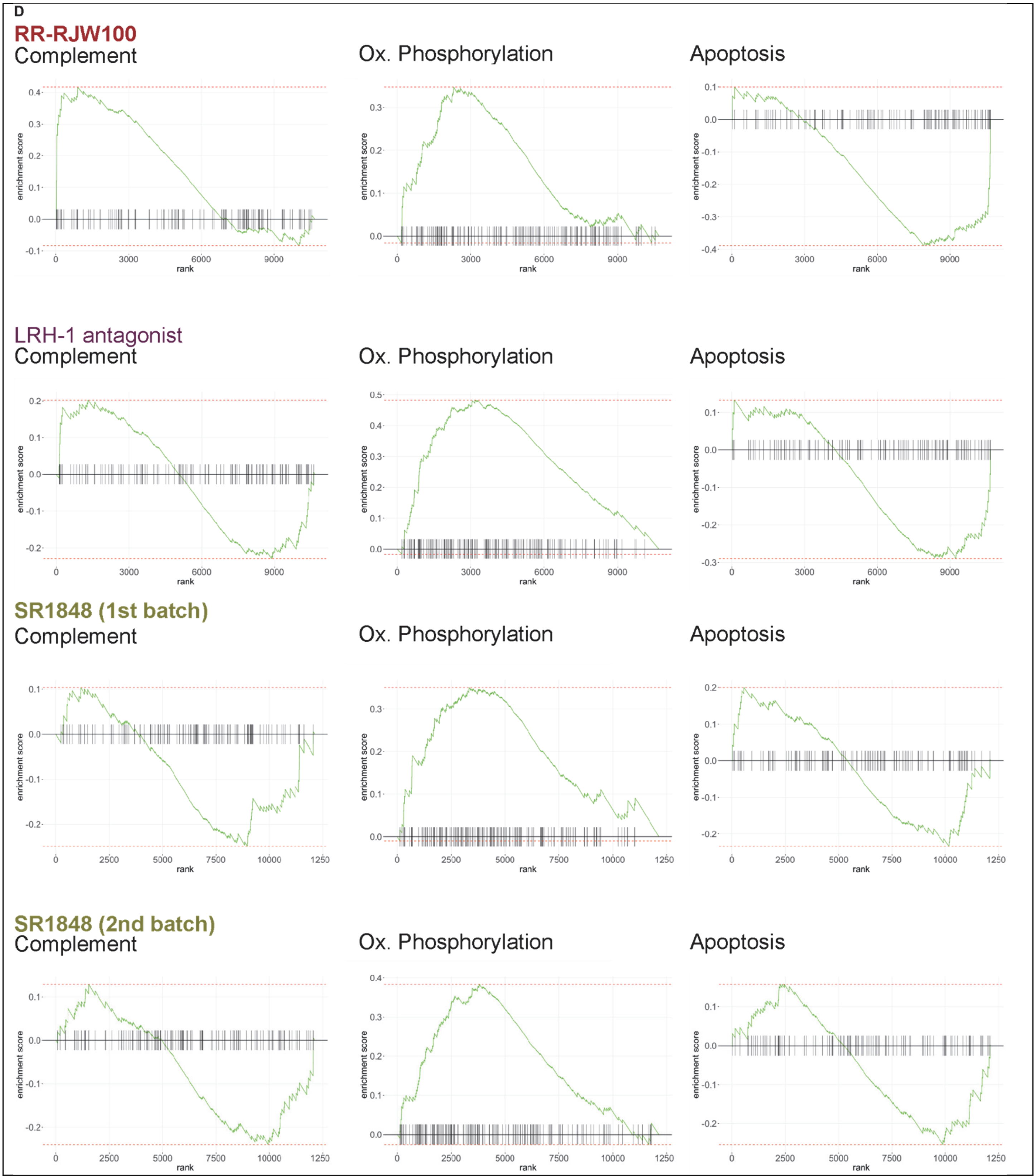


**Supplementary Figure 3.** ***Small molecule modulators of LRH-1 result in unique transcriptional responses in neutrophils***. These data correspond to those illustrated in **Figure 3** of the main text. (**A**) GO analyses for treatment of neutrophils with indicated LRH-1 ligands. BP: biological process, MF: molecular function, CC cellular component. (**B**) GO analyses for only those genes that were uniquely regulated by RR-RJW100. (**C i-ii**) GO analyses for only those genes that were commonly regulated by LRH-1 antagonist and SR1848. (**D**) Gene set enrichment analysis (GSEA), focusing on the hallmarks for the complement system, inflammatory response, oxidative phosphorylation and reactive oxygen species pathways.

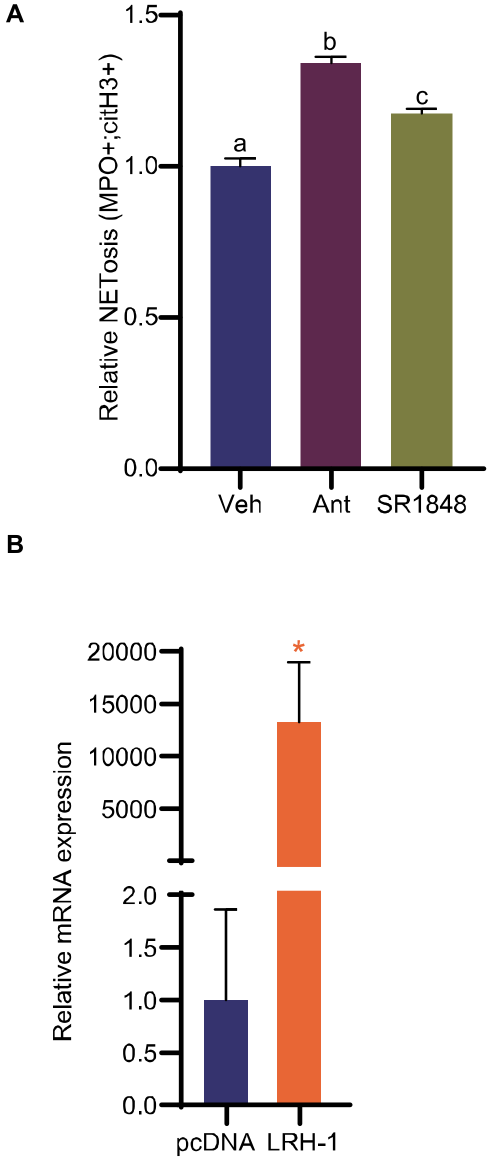


**Supplementary Figure 4.** ***LRH-1 antagonist or inverse agonist increase NETosis as assessed by double-positive extracellular staining for MPO and citH3. LRH-1 can be acutely overexpressed in neutrophils.*** (**A**) Neutrophils were treated as in **Fig. 6A**. Resulting culture was then stained form MPO and citH3 followed by flow cytometry analysis. Events were normalized and expressed as relative to vehicle control. Different letters denote statistical significance ($P<0.05$, one-way ANOVA followed by multiple comparison test with Šidáks correction). (**B**) Neutrophils that were electroporated with an LRH-1 expression plasmid had increased mRNA expression of LRH-1 (data was LN transformed for normality, unpaired T test, $P<0.05$). 1μg of pcDNA control or LRH-1 expression plasmid were nucleofected.

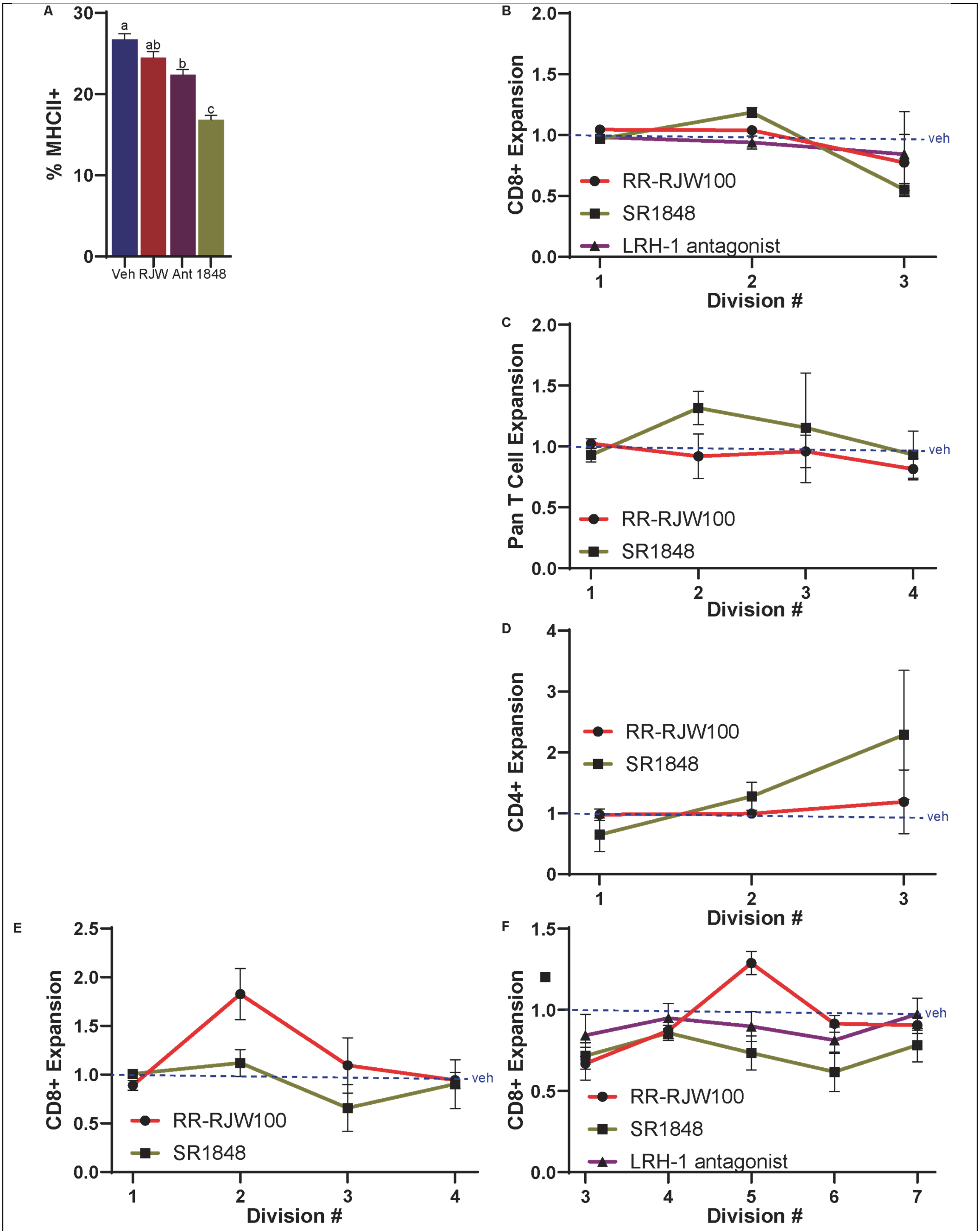


**Supplementary Figure 5.** ***Treatment of neutrophils with small molecule modulators of LRH-1 results in altered T cell expansion.*** (**A**) Neutrophils treated with LRH-1 antagonist (Ant) or SR1848 (1848) had higher expression of MHCII compared to vehicle or RR-RJW100, as determined by flow cytometry. Bone marrow derived neutrophils were cultured with LRH-1 ligands for 24 hours, then primed for 1 hour in 4T1-conditioned media. Cells were stained for surface MHC II (I-A/I-E). Different letters denote statistical significance ($P<0.05$, One-way ANVOA followed by a post-hoc test with the Šidák's correction for multiple comparisons). (**B**) Expansion of CD8+ T cells from OTII mice co-cultured with neutrophils pre-treated with the indicated ligands, OVA and LPS. (**C**) Pan-T cell expansion of OTI T cells in the presence of neutrophils pre-treated as in (A). (**D**) CD4+ specific expansion of OTI T cells in the presence of neutrophils pre-treated as in (A). (**E**) CD8+ specific expansion of T cells isolated from OTI mice, cultured with neutrophils. (**F**) CD8+ T cell expansion when pan-T cells were activated with antibodies against CD3 and CD28, as in **Fig. 7D**.

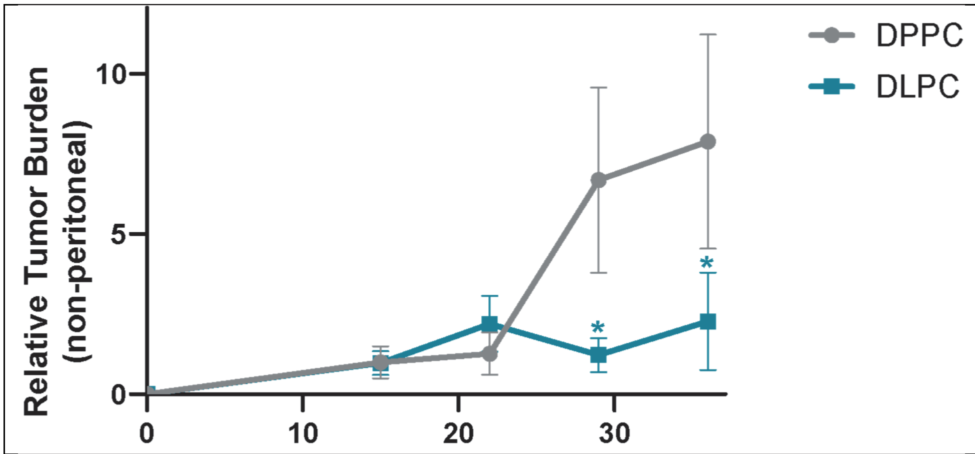


**Supplementary Figure 6.** ***LRH-1 agonist reduces metastatic burden of ID8 ovarian tumor.*** Data corresponds to **Figure 8A**. Bioluminescent imaging was performed with regions of interest placed from the diaphragm towards the rostral area. Asterisks (*) denote statistical difference between DPPC control and DLPC for the indicated timepoint (data fit to a mixed-effects model followed by a post-hoc test with the Šidák's correction for multiple comparisons).
